# Distinct patterns of *de novo* coding variants contribute to Tourette Syndrome etiology

**DOI:** 10.1101/2025.11.04.686570

**Authors:** Lingyu Zhan, Dongmei Yu, Franjo Ivankovic, Maria Niarchou, Laura Domenech-Salgado, Cathy L. Barr, Fortu Benarroch, Cathy L. Budman, Danielle C. Cath, Nelson B. Freimer, Helena Garrido, Marco A. Grados, Varda Gross-Tsur, Luis Herrera-Amighetti, Robert A. King, Roger Kurlan, James F. Leckman, Gholson J. Lyon, William M. McMahon, David L. Pauls, Yehuda Pollak, Mary M. Robertson, Roxana Romero, Guy A. Rouleau, Paul Sandor, Harvey S. Singer, Paola Giusti-Rodriguez, Lea K. Davis, Carol A. Mathews, Jeremiah M. Scharf, Roel A. Ophoff

## Abstract

Tourette syndrome (TS) is a highly heritable childhood-onset neuropsychiatric disorder characterized by persistent motor and vocal tics. While both common and rare variants contribute to TS susceptibility, the role of rare *de novo* mutations (DNMs) remains incompletely characterized. Here, we report findings from the largest TS whole-exome sequencing study to date, analyzing 1,466 TS trios alongside 6,714 autism spectrum disorder (ASD) trios and 5,880 unaffected sibling controls from the Simons Simplex Collection (SSC) and SPARK cohorts. Leveraging a trio-based design across these cohorts enabled calibrated assessment of DNM burden while controlling for background mutation rates. We observed a significant exome-wide enrichment of protein-truncating DNMs in TS probands, particularly within genes intolerant to loss-of-function variation (pLI ≥ 0.9), with little contribution from damaging missense variants. Notably, TS probands did not exhibit enrichment in previously implicated ASD or developmental delay (DD) genes, but elsewhere in the genome, suggesting a distinct rare variant architecture. Using a Bayesian statistical framework that integrates both *de novo* and rare inherited coding variants, we identified three candidate TS risk genes with FDR ≤ 0.05: *PPP5C*, *EXOC1*, and *GXYLT1*. Literature shows that they have prior links to neurodevelopmental and psychiatric disorders. These findings reveal a rare variant burden in TS that is genetically distinguishable from ASD, underscore the importance of loss-of-function mutations in TS risk, and nominate novel candidate genes for future functional investigation.

## Introduction

Tourette syndrome (TS) is a childhood-onset developmental neuropsychiatric disorder that is characterized by the presence of multiple motor and vocal tics that wax and wane and persist for at least one year.(1) TS affects approximately 0.6-0.8% of children and adults worldwide, and recent prevalence studies conducted by the Centers for Disease Control estimate that 350,000 — 450,000 individuals in the US are currently living with TS.(2–4) Furthermore, persistent motor or vocal tic disorder (PMVT), previously known as chronic tic disorder prior to DSM-5 (5), is present in an additional 1.3-1.6% of children and adolescents. (2,3,6) In part due to evidence suggesting shared genetic etiologies, TS and PMVT are now considered to lie on a spectrum or continuum of tic disorders.(6,7) In addition to impairment and distress caused directly by tics themselves, individuals with TS and PMVT have high rates of multiple, childhood-onset neuropsychiatric and neurodevelopmental disorders (NDDs), including obsessive compulsive disorder (OCD), attention deficit hyperactivity disorder (ADHD), and autism spectrum disorder (ASD)(8,9) These disorders share common features with TS (e.g., early age of onset, restrictive, repetitive, or impulsive/disinhibited behaviors that interrupt intended cognitive, motor, behavioral and social function). Genetic studies have increasingly also suggested shared underlying genetic susceptibility.(10)

TS is among the most heritable of the NDDs, with an estimated population-based heritability of 70-80%, within which 58% is directly attributable to SNPs.(11,12) As a comparison, population-based heritability estimates for OCD, ADHD, and ASD are 25-48% (12–16), 60-90% (17,18), and 50-90% (19,20), respectively. TS is also highly polygenic, with both common and rare variants contributing to TS genetic risk.(12,21) Although requiring replication, genome-wide association studies (GWAS) have identified multiple genome-wide significant loci potentially associated with TS, including *BCL11B, NDFIP2, RBM26, MCHR2-AS1, DRAM1, COL27A1, NR2F1, and FLT3*.(7,22–25)

Recent studies have also demonstrated that rare, gene-damaging variants (RVs) contribute to genetic risk for many neuropsychiatric disorders, such as ASD, schizophrenia (SCZ), and bipolar disorder (BP).(26–32) Of particular interest, RVs contribute a substantial proportion of genetic risk to OCD and ASD, both of which, as noted, have significant clinical and genetic overlap with TS. Although fewer studies have been conducted, analyses of RVs have also identified additional potential TS risk genes, including *CNTN6*, *NRXN1*, *SLITRK1*, *HDC*, and *CELSR3*, highlighting the importance of the investigation of the rare spectrum of genetic variation.(7,21,33)

A key interest in studies on TS is to explore the connection between TS and ASD, which demonstrate considerable clinical and etiological overlap. Epidemiological studies reveal a higher-than-expected co-occurrence of these two disorders, with a notable percentage of individuals with TS exhibiting autistic symptoms or meeting diagnostic criteria for ASD. (8,34,35) This symptomatic convergence includes features such as repetitive behaviors (manifesting as tics in TS and stereotypies in ASD), echolalia, and challenges in social communication. Accumulating research indicates shared genetic susceptibilities, with rare copy number variants (CNVs) and disruptions in genes like *CNTNAP2*, *NRXNs*, and *NLGNs*, as well as common variants identified from GWAS studies implicated in both disorders (34,36–38). These shared genetic factors point towards common pathophysiological mechanisms, potentially involving dysregulation of cortico-striatal-thalamo-cortical (CSTC) circuitry and broader neurodevelopmental pathways (34,39). However, the shared genetic basis between TS and ASD, especially the contribution of RVs, is understudied, and the TS-specific risk genes are not fully explored.

Large-scale sequencing data are typically required to achieve adequate statistical power in rare-variant research. Here, we present the results of the largest TS whole-exome sequencing (WES) study to date, with 1,466 TS trio families (parent-proband trios) available for analysis. A subset of this sample has previously been analyzed and yielded a number of potential TS risk genes (*CNTN6*, *NRXN1*).(40) The current analysis also includes 2,483 sib-pair families from the Simons Simplex Cohort (SSC) and 4,231 sib-pair families from the Simons Powering Autism Research (SPARK) cohorts (e.g., families with one ASD proband and one unaffected sibling). These samples are included in the analysis to serve as: 1) unaffected controls (unaffected siblings from the SSC and SPARK ASD families) and 2) clinical controls (ASD-affected probands from SSC and SPARK families). Leveraging the unique pedigree structure of these three cohorts, we investigated the effects of *de novo mutation* (DNM) burden on TS both at the exome-wide level and in sets of genes, and compared them to those in ASD. Gene-specific analysis was conducted using a Bayesian statistical framework (e.g., transmission and *de novo* association; TADA), to incorporate the effects of both inherited and *de novo* gene-damaging variant associations with TS, resulting in the identification of three novel TS risk genes.(41,42) Overall, our large sample size and ability to incorporate family-level data into the analyses provide new insights into the contribution of rare coding variants in TS and new perspectives on our understanding of the underlying genetic etiology of TS. An overview of the study workflow is summarized in Figure 1.

**Figure 1.**
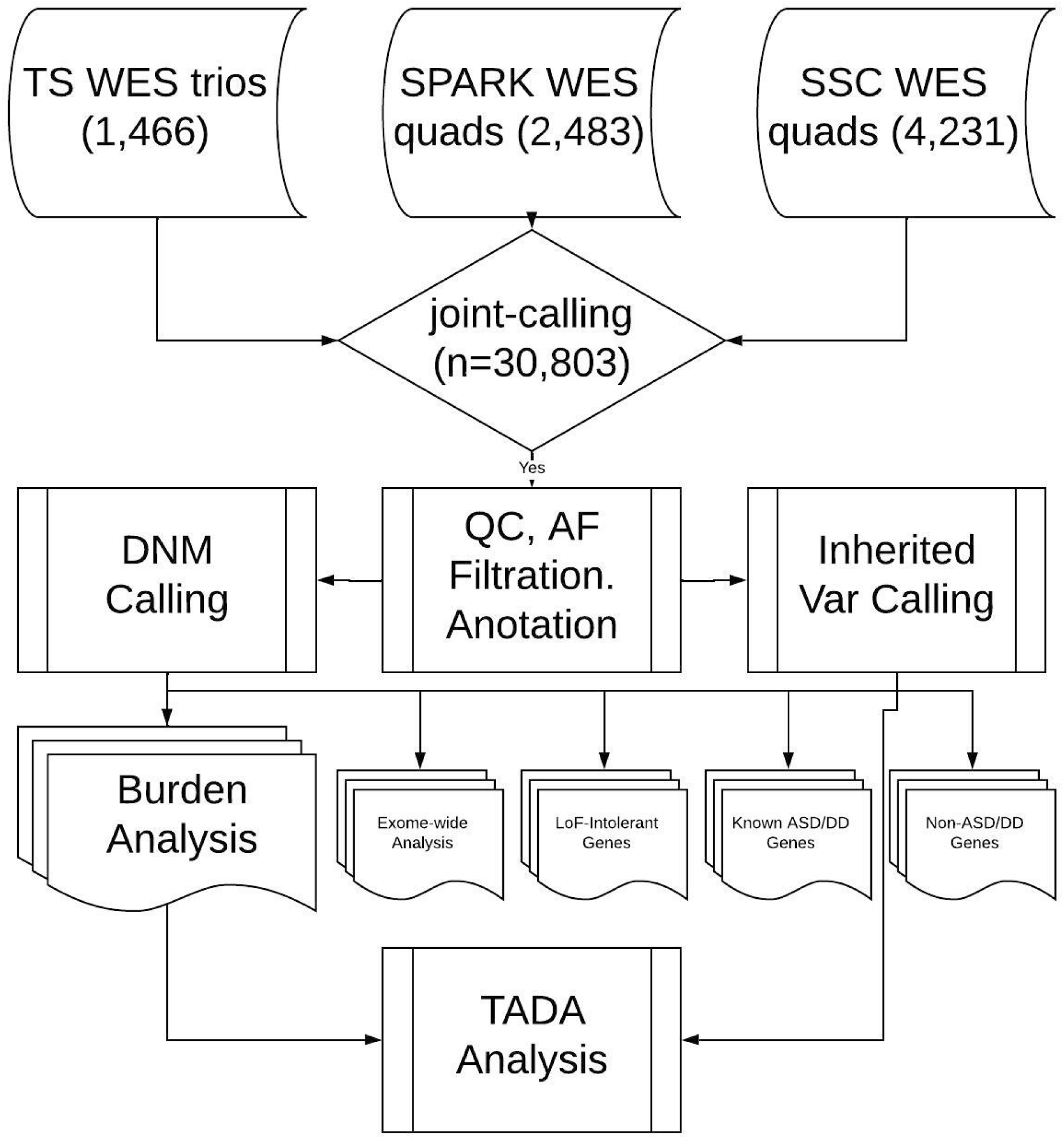
Diagram of the study.

## Results

### Discovery of rare and *de novo* coding variants in TS and ASD cohorts

We aggregated and jointly called variants from exome sequencing data across three cohorts: 4,430 individuals from the TS cohort, 9,449 individuals from the SSC cohort, and 16,924 individuals from the SPARK cohort. Due to differences in sequencing platforms, we utilized the overlapping exome regions and applied stringent quality controls to control for batch effects. We retained a total of 3,950 individuals from the TS cohort, consisting of 1,413 affected probands, seven unaffected children, and 2,530 parents, of which 65.14% of the exome data has not been previously published. From the SSC and SPARK cohorts, we retained 8,458 and 16,920 individuals, respectively, comprising 6,744 ASD probands, 5,880 unaffected siblings, and 12,754 parents. We then applied stringent variant quality filterings and subsequently selected rare variants (MAC < 20) using the non-neuro subset of the gnomAD exome database as the reference panel.(43) After calling *de novo* mutations using TrioDeNovo and removing low-quality candidates, we obtained an average of 1.28 *de novo* single-nucleotide variants (SNVs) per individual, with a rate of 1.54 in TS probands, 1.27 in ASD cases from SSC, and 1.26 in ASD cases from SPARK. (Table 1)

**Table 1.** Rate of DNMs in each cohort, split by variant annotations. Syn, synonymous variants; PTV, protein-truncating variants; Dmis, missense variants with a REVEL score ≥ 0.644.

| Cohort | All DNMs | Syn | PTV | Dmis |
| --- | --- | --- | --- | --- |
| TS cases | 0.906 | 0.207 | 0.0505 | 0.0822 |
| SSC cases | 0.842 | 0.189 | 0.0537 | 0.0835 |
| SSC controls | 0.771 | 0.194 | 0.0305 | 0.0707 |
| SPARK cases | 0.871 | 0.195 | 0.0530 | 0.0956 |
| SPARK controls | 0.864 | 0.208 | 0.0340 | 0.0749 |

### An exome-wide enrichment of pathogenic DNMs was observed in TS probands

In the first stage of analysis, we sought to explore the overall pattern of *de novo* coding variants in both TS and ASD cohorts. Two categories of pathogenic variants were used in the analysis: protein-truncating variants (PTVs) and damaging missense variants (Dmis) (See Methods). Following existing recommendations, the pathogenicity of missense variants was annotated with three REVEL score thresholds: ≥0.644 (most lenient), ≥0.773, and ≥0.932 (most stringent).(44,45) Attempting to be inclusive, in this study, we focused on the first (most lenient) category, which encompassed the largest number of Dmis variants. Using unaffected siblings from the SSC and SPARK cohorts as controls, we observed significant exome-wide enrichment of pathogenic DNMs (including PTVs and all Dmis variants) in the TS probands compared to unaffected control siblings (OR = 1.23∼1.32, P-value = 9.69e-3∼2.21e-2). (Figure 2 and Table 2) Similarly, significant burdens of pathogenic DNMs were observed in ASD probands from the SSC (OR = 1.26∼1.36, P-value = 1.54e-3∼1.79e-3) and SPARK (OR = 1.36∼1.47, P-value = 3.49e-7∼9.11e-6) cohorts compared to unaffected siblings. The observed enrichment was further elevated in both TS and ASD probands when we focused solely on PTVs (TS-control: OR = 1.49∼1.67, P-value = 5.78e-3∼6.55e-3; ASD-control: OR = 1.57∼1.78, P-value = 2.50e-5∼6.59e-4). (Figure 2 and Table 2) While the Dmis variants were significantly enriched in ASD probands in both the SSC and the SPARK cohort, we did not observe a statistically significant elevation of Dmis variants in the TS probands, suggesting that PTVs were the main contributor to the exome-wide burden in TS. Analysis results using Dmis variants from the other two categories were included in Supplementary Figures 1 and 2.

**Figure 2.**
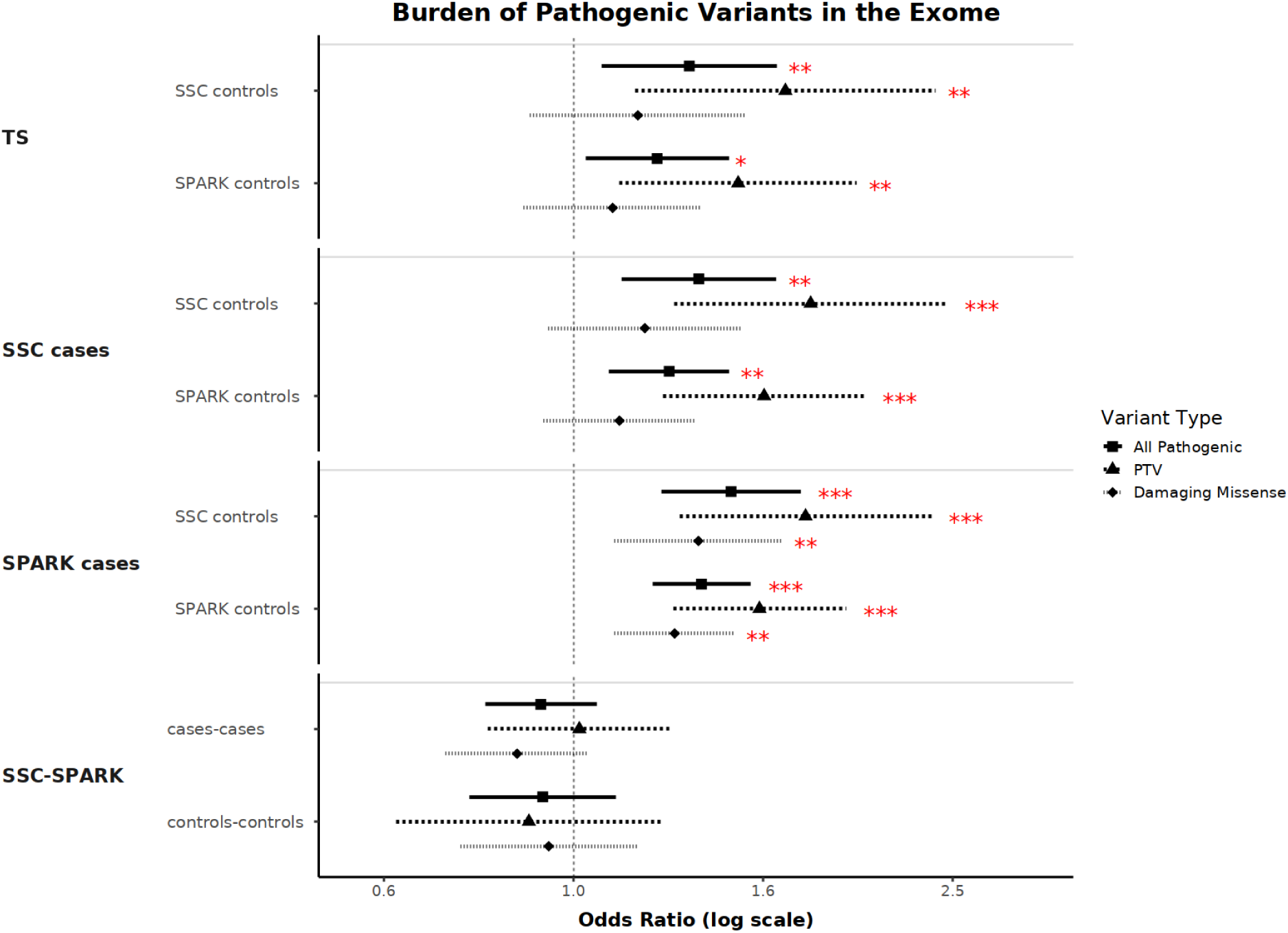
Forest plot of the burden analysis of DNMs between cohorts within the context of the exome, split by variant annotations. All pathogenic, PTV + Dmis; PTV, protein-truncating variants; Dmis, missense variants with a REVEL score ≥ 0.644.

**Table 2.** Burden analysis of DNMs between cohorts within the context of the exome, split by variant annotations. All pathogenic, PTV + Dmis; PTV, protein-truncating variants; Dmis, missense variants with a REVEL score ≥ 0.644.

| Comparison |  | All Pathogenic |  | PTV |  | Dmis |  |
| --- | --- | --- | --- | --- | --- | --- | --- |
| Sample 1 | Sample 2 | OR | Pvalue | OR | Pvalue | OR | Pvalue |
| TS cases | SSC Controls | 1.32 | 9.69E-03 | 1.67 | 5.78E-03 | 1.17 | 2.42E-01 |
|  | SPARK Controls | 1.23 | 2.21E-02 | 1.49 | 6.55E-03 | 1.10 | 3.91E-01 |
| SSC cases | SSC Controls | 1.36 | 1.54E-03 | 1.78 | 6.59E-04 | 1.19 | 1.49E-01 |
|  | SPARK controls | 1.26 | 1.79E-03 | 1.59 | 2.18E-04 | 1.12 | 2.36E-01 |
| SPARK cases | SSC Controls | 1.47 | 9.11E-06 | 1.76 | 3.02E-04 | 1.35 | 3.59E-03 |
|  | SPARK controls | 1.36 | 3.49E-07 | 1.57 | 2.50E-05 | 1.28 | 1.00E-03 |
| SSC-SPARK | Cases-cases | 0.92 | 2.51E-01 | 1.01 | 9.01E-01 | 0.87 | 1.21E-01 |
|  | Controls-controls | 0.93 | 4.11E-01 | 0.90 | 5.10E-01 | 0.94 | 5.83E-01 |

### The observed *de novo* burden is concentrated in LoF-intolerant genes

Next, we focused our analysis on LoF-intolerant genes, which are considered to impact essential human fitness.(46) We applied two separate criteria: 1) pLI score ≥ 0.9, 2) LOEUF < 0.6, which produced 3,231 and 3,784 LoF-intolerant genes, respectively, with 2,366 genes in overlap. (See Methods) Using LoF-intolerant genes based on the pLI metric, we observed an excess of pathogenic, gene-damaging *de novo* SNVs in TS probands compared with unaffected siblings from the SSC and SPARK cohorts (OR = 1.56∼1.58, P-value = 3.50e-3∼1.50e-2), as well as in ASD probands from the SSC (OR = 1.96∼1.98, P-value = 9.10e-8∼4.43e-5) and SPARK cohorts (OR = 2.30∼2.34, P-value = 1.09e-14∼4.50e-8). (Figure 3 and Table 3) The effect sizes of pathogenic DNMs in both TS and ASD were more elevated in LoF-intolerant genes than the overall exome-wide scale, suggesting that DNMs observed in both disorders primarily affect core human functions. We next looked at the distribution of PTVs and Dmis variants in the LoF-intolerant genes. This analysis demonstrated a significant concentration of DNM burden of PTVs within the LoF-intolerant genes for both the TS probands (OR = 2.21∼2.66, P-value = 3.29e-3∼7.61e-3) and the ASD probands (OR = 2.84∼3.77, P-value = 1.08e-8∼2.47e-4). (Figure 3 and Table 3) The Dmis variants showed a trend for enrichment in the TS probands while reaching statistical significance in the ASD probands from both the SSC and SPARK cohorts, though with smaller effect sizes than the PTVs, suggesting that PTVs explained most of the DNM burden in both disorders.

**Figure 3.**
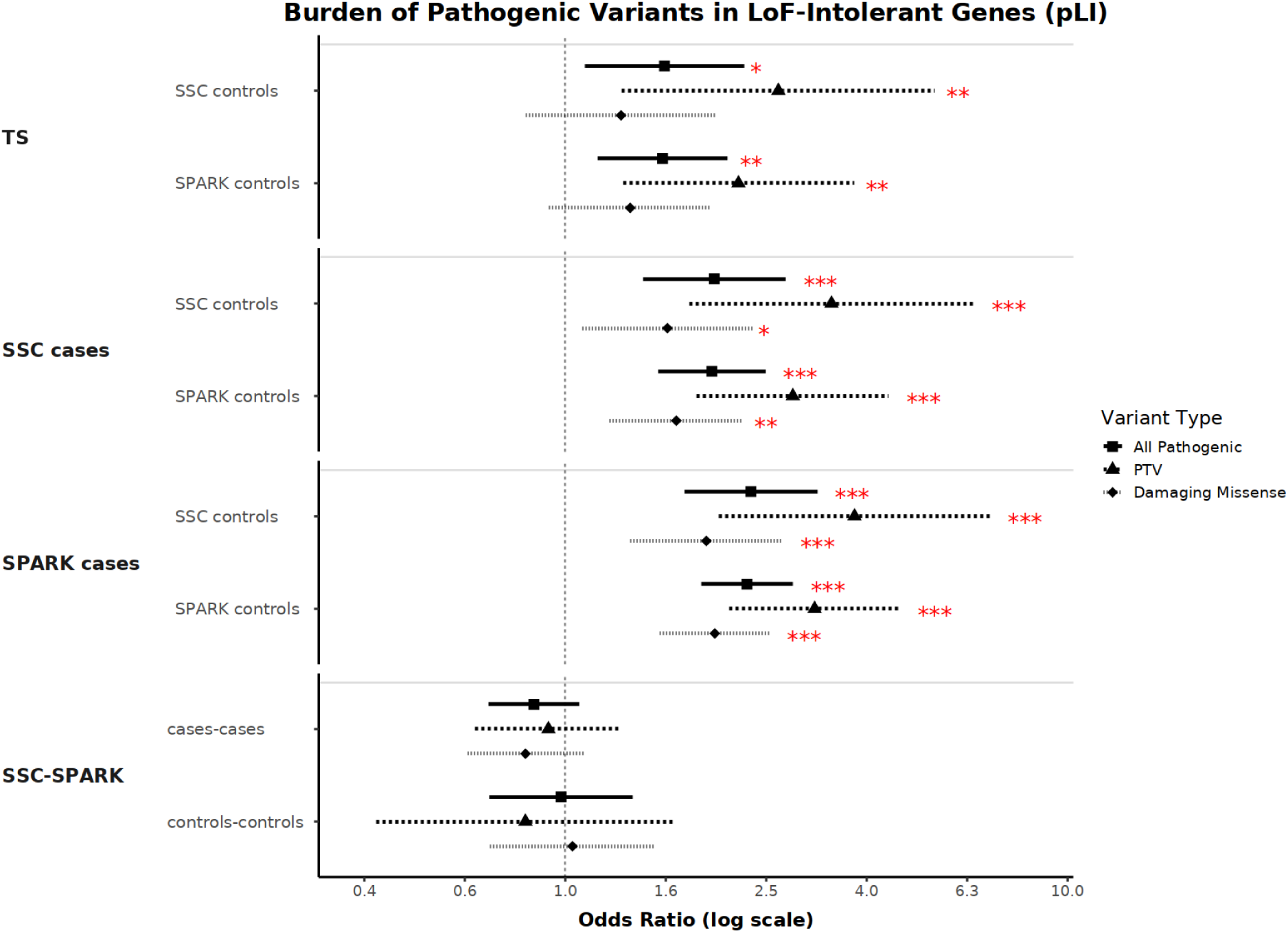
Forest plot of the burden analysis of DNMs between cohorts within the context of the LoF-intolerant genes (defined by pLI score), split by variant annotations. All pathogenic, PTV + Dmis; PTV, protein-truncating variants; Dmis, missense variants with a REVEL score ≥ 0.644.

**Table 3.** Burden analysis of DNMs between cohorts within the context of the LoF-intolerant genes (defined by pLI score), split by variant annotations. All pathogenic, PTV + Dmis; PTV, protein-truncating variants; Dmis, missense variants with a REVEL score ≥ 0.644.

| Comparison |  | All Pathogenic |  | PTV |  | Dmis |  |
| --- | --- | --- | --- | --- | --- | --- | --- |
| Sample 1 | Sample 2 | OR | Pvalue | OR | Pvalue | OR | Pvalue |
| TS | SSC Controls | 1.58 | 1.50E-02 | 2.66 | 7.61E-03 | 1.29 | 2.51E-01 |
|  | SPARK Controls | 1.56 | 3.50E-03 | 2.21 | 3.29E-03 | 1.35 | 1.15E-01 |
| SSC cases | SSC Controls | 1.98 | 4.43E-05 | 3.39 | 2.47E-04 | 1.60 | 1.84E-02 |
|  | SPARK controls | 1.96 | 9.10E-08 | 2.84 | 3.40E-06 | 1.66 | 1.04E-03 |
| SPARK cases | SSC Controls | 2.34 | 4.50E-08 | 3.77 | 2.99E-05 | 1.91 | 2.91E-04 |
|  | SPARK controls | 2.30 | 1.09E-14 | 3.14 | 1.08E-08 | 1.98 | 8.49E-08 |
| SSC-SPARK | Cases-cases | 0.87 | 1.76E-01 | 0.93 | 6.52E-01 | 0.83 | 1.75E-01 |
|  | Controls-controls | 0.98 | 9.09E-01 | 0.83 | 6.01E-01 | 1.03 | 8.64E-01 |

With LoF-intolerant genes based on LOEUF score, we observed a similar enrichment of pathogenic variants in TS probands (P-value = 2.46e-2∼2.51e-2), SSC ASD probands (P-value = 1.33e-6∼5.11e-5), and SPARK ASD probands (P-value = 1.03e-10∼5.62e-7), though the ORs were lower than those estimated from the pLI-defined genes in all comparisons. (Figure 4 and Table 4) The PTVs showed a similar increase in ORs (TS-control: OR = 1.81∼2.00, P-value = 1.36e-2∼2.48e-2; ASD-control: OR = 2.30∼2.84, P-value = 9.54e-7∼3.62e-4), validating our previous results. Analysis, including missense variants of different REVEL score thresholds, can be found in Supplementary Figures 3-6.

**Figure 4.**
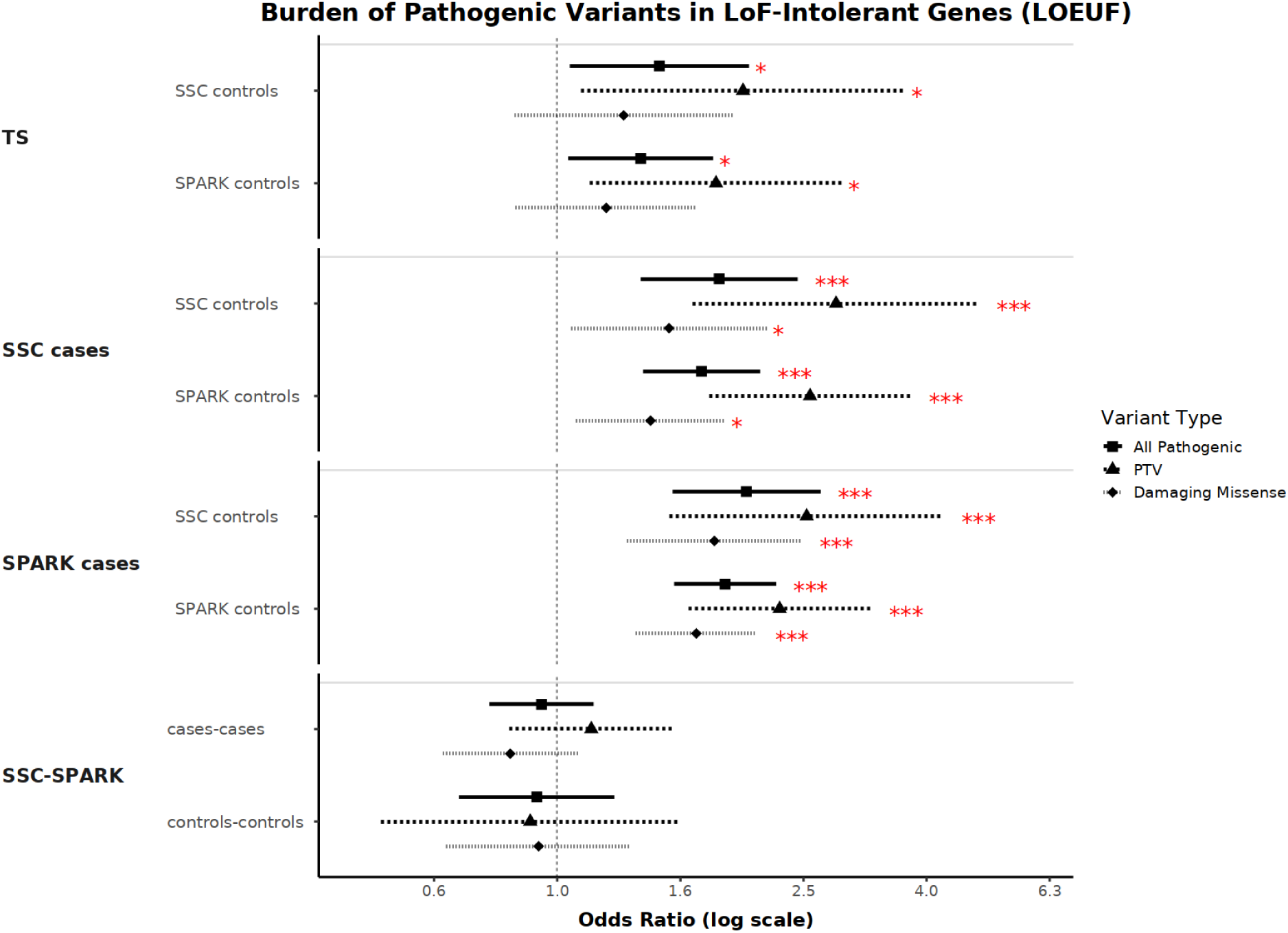
Forest plot of the burden analysis of DNMs between cohorts within the context of the LoF-intolerant genes (defined by LOEUF metric), split by variant annotations. All pathogenic, PTV + Dmis; PTV, protein-truncating variants; Dmis, missense variants with a REVEL score ≥ 0.644.

**Table 4.** Burden analysis of DNMs between cohorts within the context of the LoF-intolerant genes (defined by LOEUF metric), split by variant annotations. All pathogenic, PTV + Dmis; PTV, protein-truncating variants; Dmis, missense variants with a REVEL score ≥ 0.644.

| Comparison |  | All Pathogenic |  | PTV |  | Dmis |  |
| --- | --- | --- | --- | --- | --- | --- | --- |
| Sample 1 | Sample 2 | OR | Pvalue | OR | Pvalue | OR | Pvalue |
| TS | SSC Controls | 1.47 | 2.51E-02 | 2.00 | 2.48E-02 | 1.28 | 2.34E-01 |
|  | SPARK Controls | 1.37 | 2.46E-02 | 1.81 | 1.36E-02 | 1.20 | 2.90E-01 |
| SSC cases | SSC Controls | 1.83 | 5.11E-05 | 2.84 | 1.37E-04 | 1.52 | 2.49E-02 |
|  | SPARK controls | 1.72 | 1.33E-06 | 2.57 | 9.54E-07 | 1.42 | 1.44E-02 |
| SPARK cases | SSC Controls | 2.03 | 5.62E-07 | 2.54 | 3.62E-04 | 1.80 | 4.18E-04 |
|  | SPARK controls | 1.87 | 1.03E-10 | 2.30 | 1.92E-06 | 1.68 | 7.20E-06 |
| SSC-SPARK | Cases-cases | 0.94 | 5.56E-01 | 1.14 | 4.18E-01 | 0.84 | 1.71E-01 |
|  | Controls-controls | 0.93 | 6.07E-01 | 0.90 | 7.23E-01 | 0.93 | 6.94E-01 |

### Known ASD and DD genes did not explain the DNM burden in TS

To explore the shared genetic architecture between TS, ASD, and DD, we identified previously established ASD (n=185)(47) and DD (n=285)(48) risk genes and characterized the distribution of DNMs from this set in our sample. As expected, the pathogenic DNMs in ASD risk genes demonstrated significant enrichment in ASD cases from both the SSC and SPARK cohorts (OR = 6.93∼11.63, P-value = 1.29e-12∼4.33e-6) compared to the unaffected siblings, recapitulating the importance of these known ASD risk genes from previous studies (Figure 5 and Table 5). A similar enrichment of pathogenic DNMs was also observed for DD genes (OR = 4.24∼4.91, P-value = 3.22e-11∼9.32e-6), present in ASD cases, though with smaller effect sizes, implicating a shared genetic effect between ASD and DD,. (Figure 6 and Table 6) However, when we examined the TS probands, the fold enrichment of pathogenic DNMs from either known ASD or DD genes was greatly attenuated and did not reach statistical significance.

**Figure 5.**
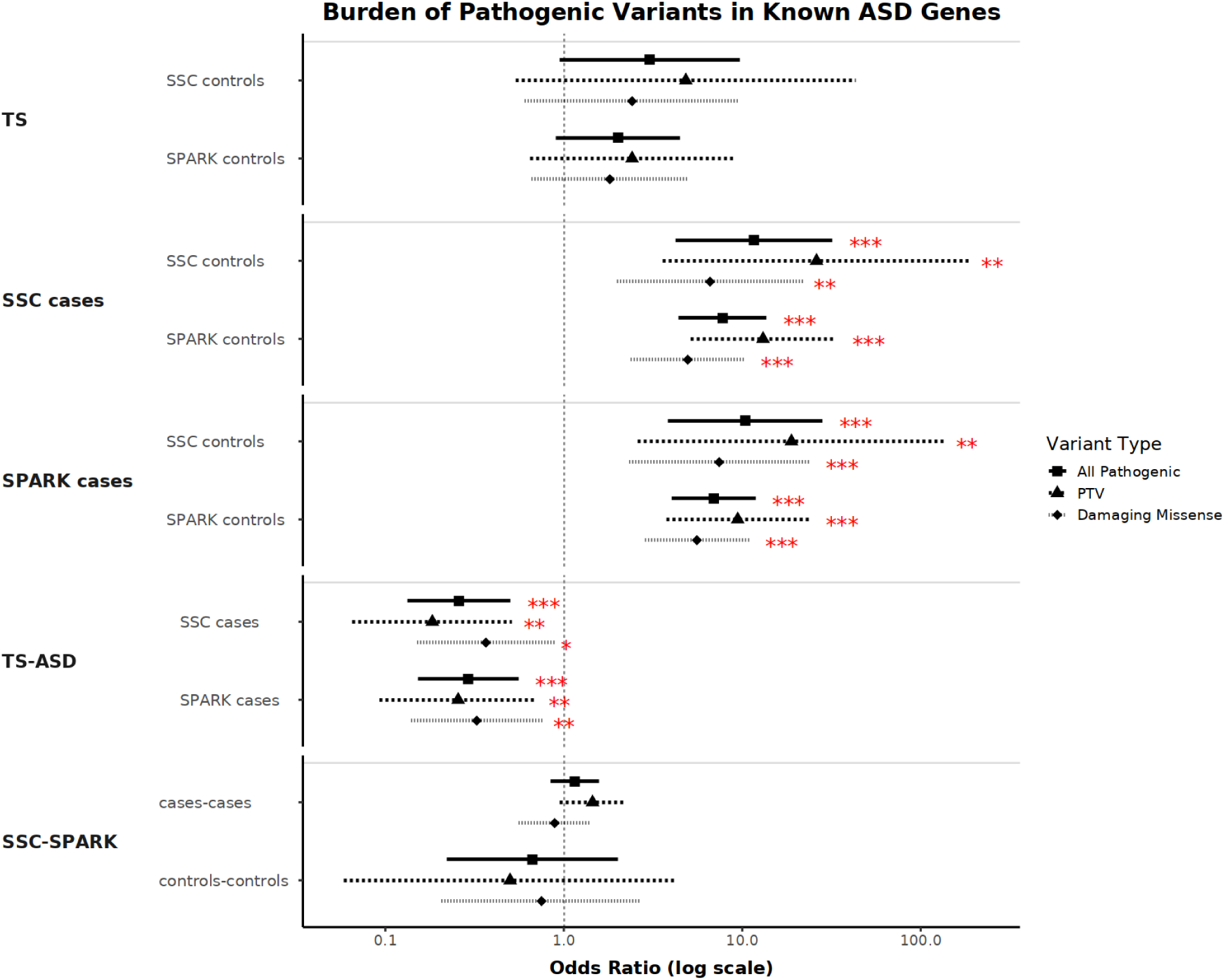
Forest plot of the burden analysis of DNMs between cohorts within the context of the known ASD genes, split by variant annotations. All pathogenic, PTV + Dmis; PTV, protein-truncating variants; Dmis, missense variants with a REVEL score ≥ 0.644.

**Figure 6.**
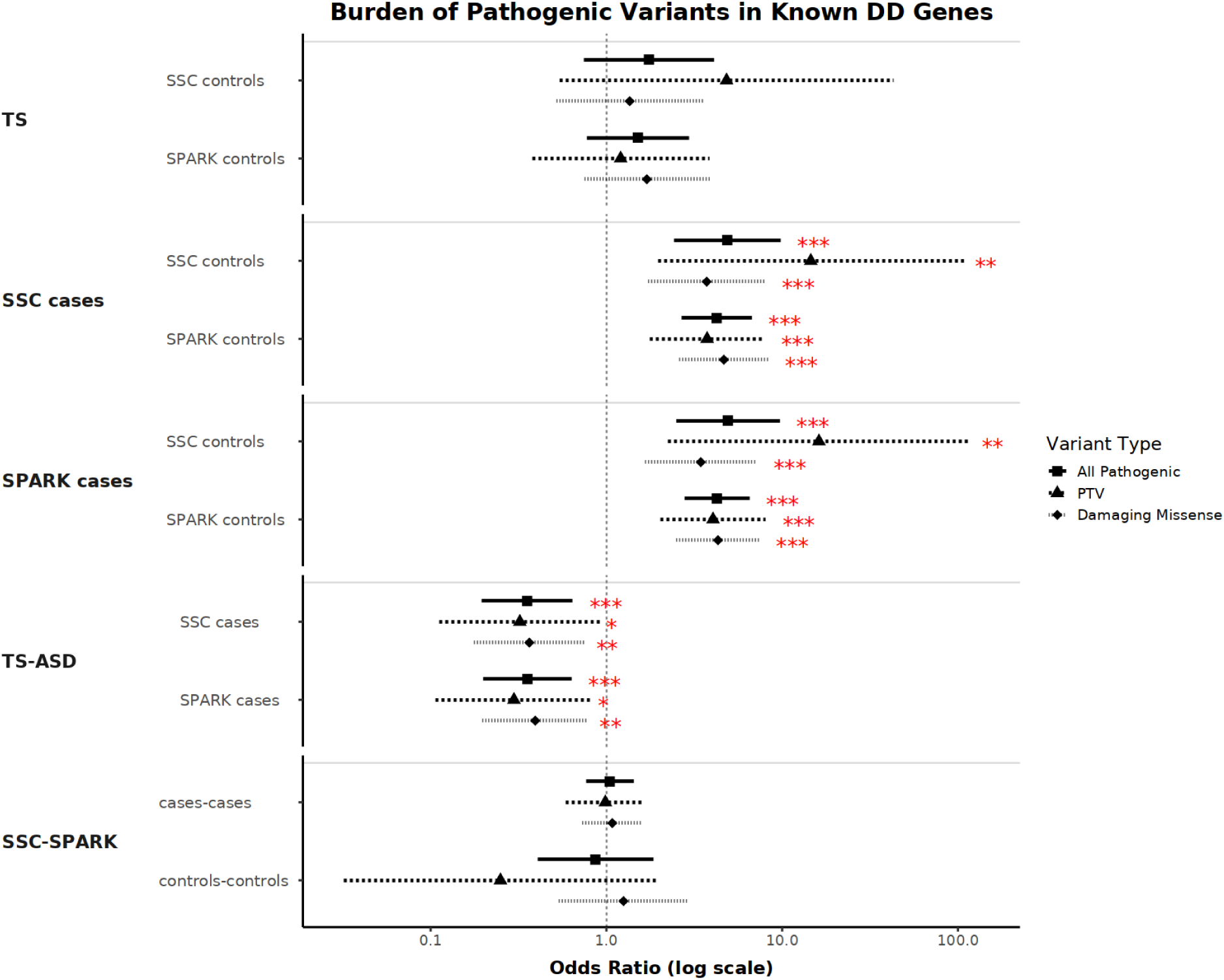
Forest plot of the burden analysis of DNMs between cohorts within the context of the known DD genes, split by variant annotations. All pathogenic, PTV + Dmis; PTV, protein-truncating variants; Dmis, missense variants with a REVEL score ≥ 0.644.

**Table 5.** Burden analysis of DNMs between cohorts within the context of the known ASD genes, split by variant annotations. All pathogenic, PTV + Dmis; PTV, protein-truncating variants; Dmis, missense variants with a REVEL score ≥ 0.644.

| Comparison |  | All Pathogenic |  | PTV |  | Dmis |  |
| --- | --- | --- | --- | --- | --- | --- | --- |
| Sample 1 | Sample 2 | OR | Pvalue | OR | Pvalue | OR | Pvalue |
| TS | SSC Controls | 3.02 | 6.19E-02 | 4.83 | 1.59E-01 | 2.41 | 2.13E-01 |
|  | SPARK Controls | 2.02 | 8.69E-02 | 2.41 | 1.89E-01 | 1.81 | 2.51E-01 |
| SSC cases | SSC Controls | 11.63 | 2.03E-06 | 26.08 | 1.26E-03 | 6.60 | 2.00E-03 |
|  | SPARK Controls | 7.77 | 1.29E-12 | 13.08 | 7.60E-08 | 4.95 | 1.81E-05 |
| SPARK cases | SSC Controls | 10.40 | 4.33E-06 | 18.86 | 3.65E-03 | 7.42 | 7.16E-04 |
|  | SPARK Controls | 6.93 | 2.43E-12 | 9.43 | 1.75E-06 | 5.57 | 5.37E-07 |
| TS | SSC Cases | 0.26 | 7.33E-05 | 0.18 | 1.26E-03 | 0.37 | 2.69E-02 |
|  | SPARK Cases | 0.29 | 1.97E-04 | 0.26 | 8.78E-03 | 0.33 | 8.92E-03 |
| SSC-SPARK | Cases-Cases | 1.15 | 3.80E-01 | 1.45 | 8.86E-02 | 0.89 | 6.18E-01 |
|  | Controls-Controls | 0.67 | 4.71E-01 | 0.50 | 5.27E-01 | 0.75 | 6.63E-01 |

**Table 6.** Burden analysis of DNMs between cohorts within the context of the known DD genes, split by variant annotations. All pathogenic, PTV + Dmis; PTV, protein-truncating variants; Dmis, missense variants with a REVEL score ≥ 0.644.

| Comparison |  | All Pathogenic |  | PTV |  | Dmis |  |
| --- | --- | --- | --- | --- | --- | --- | --- |
| Sample 1 | Sample 2 | OR | Pvalue | OR | Pvalue | OR | Pvalue |
| TS | SSC Controls | 1.75 | 2.00E-01 | 4.83 | 1.59E-01 | 1.36 | 5.31E-01 |
|  | SPARK Controls | 1.51 | 2.26E-01 | 1.21 | 7.52E-01 | 1.70 | 2.05E-01 |
| SSC cases | SSC Controls | 4.87 | 9.32E-06 | 14.55 | 8.80E-03 | 3.72 | 7.40E-04 |
|  | SPARK Controls | 4.24 | 8.25E-10 | 3.74 | 5.46E-04 | 4.66 | 2.42E-07 |
| SPARK cases | SSC Controls | 4.91 | 4.46E-06 | 16.19 | 5.93E-03 | 3.45 | 9.27E-04 |
|  | SPARK Controls | 4.25 | 3.22E-11 | 4.04 | 7.08E-05 | 4.31 | 1.29E-07 |
| TS | SSC Cases | 0.35 | 6.59E-04 | 0.32 | 3.55E-02 | 0.36 | 6.59E-03 |
|  | SPARK Cases | 0.36 | 4.37E-04 | 0.30 | 2.08E-02 | 0.39 | 8.60E-03 |
| SSC-SPARK | Cases-Cases | 1.04 | 7.84E-01 | 0.99 | 9.59E-01 | 1.08 | 7.00E-01 |
|  | Controls-Controls | 0.87 | 7.09E-01 | 0.25 | 1.86E-01 | 1.25 | 6.04E-01 |

We further examined the PTVs alone, where we observed the highest fold of enrichment in previous analyses with known ASD and DD genes. We again observed a higher enrichment of PTVs in both gene sets in ASD probands (ASD: OR = 9.43∼26.08, P-value = 7.60e-8∼3.65e-3; DD: OR = 3.74∼16.19, P-value = 7.08e-5∼8.80e-3) than those identified in the previous analyses. In contrast, a non-significant elevation of PTVs was detected in TS probands, though with a slight increase in effect sizes compared to what was seen with all pathogenic DNMs (Figures 5 and Table 5). An additional analysis using Dmis variants in known ASD and DD genes revealed that the enrichment pattern present in ASD probands and absent in TS probands, remained significant, further suggesting that both PTVs and Dmis variants within these two gene sets contribute to the development of ASD but not to TS. Analyses using missense variants with a REVEL score of 0.773 or higher are available in Supplementary Figures 7-8.

Additionally, we observed that all types of PTVs and Dmis variants examined in these two gene sets demonstrated a statistically significant depletion in TS probands, compared to ASD and DD probands (ASD: OR = 0.18∼0.37, P-value = 7.33e-5∼2.69e-2; DD: OR = 0.30∼0.39, P-value = 4.37e-4∼3.55e-2), indicating that the known ASD and DD genes conferred less risk on TS than on ASD and DD, and consequently implying that the core genes affecting TS might be different from the ones affecting ASD.

### TS and ASD probands from SPARK shared a different genetic architecture from ASD probands from SSC

Given these results from the ASD/DD gene-set analysis described above, we hypothesized that the exome-wide DNM burden in TS probands present in the first analysis should be located in other parts of the genome. To test this hypothesis, we removed ASD and DD susceptibility genes, respectively, from the analyses. As expected, when ASD risk genes were excluded, the SSC ASD probands demonstrated no significant enrichment of pathogenic DNMs or PTVs individually. In contrast, statistically significant enrichment of both pathogenic DNMs (OR = 1.20∼1.28, P-value = 2.45e-2∼4.78e-2) and PTVs alone (OR = 1.46∼1.62, P-value = 1.17e-2∼1.22e-2) remained present in the TS trio probands, suggesting a potentially distinct, underlying rare variant genetic basis. (Figure 7 and Table 7) Furthermore, when we compared the SPARK ASD probands to the unaffected siblings, we observed significant remaining elevation of pathogenic DNMs (OR = 1.18∼1.26, P-value = 9.65e-3∼9.89e-3), as well as PTVs alone (OR = 1.28∼1.41, P-value = 2.95.0e-2∼3.35e-2) in ASD probands, after the ASD genes were omitted. Given the heterogeneity of phenotypes in the SPARK cohort, this observation may imply a different genetic architecture in the SPARK ASD samples compared to the SSC ASD samples, possibly due to the contribution of other neuropsychiatric conditions present in the data.

**Figure 7.**
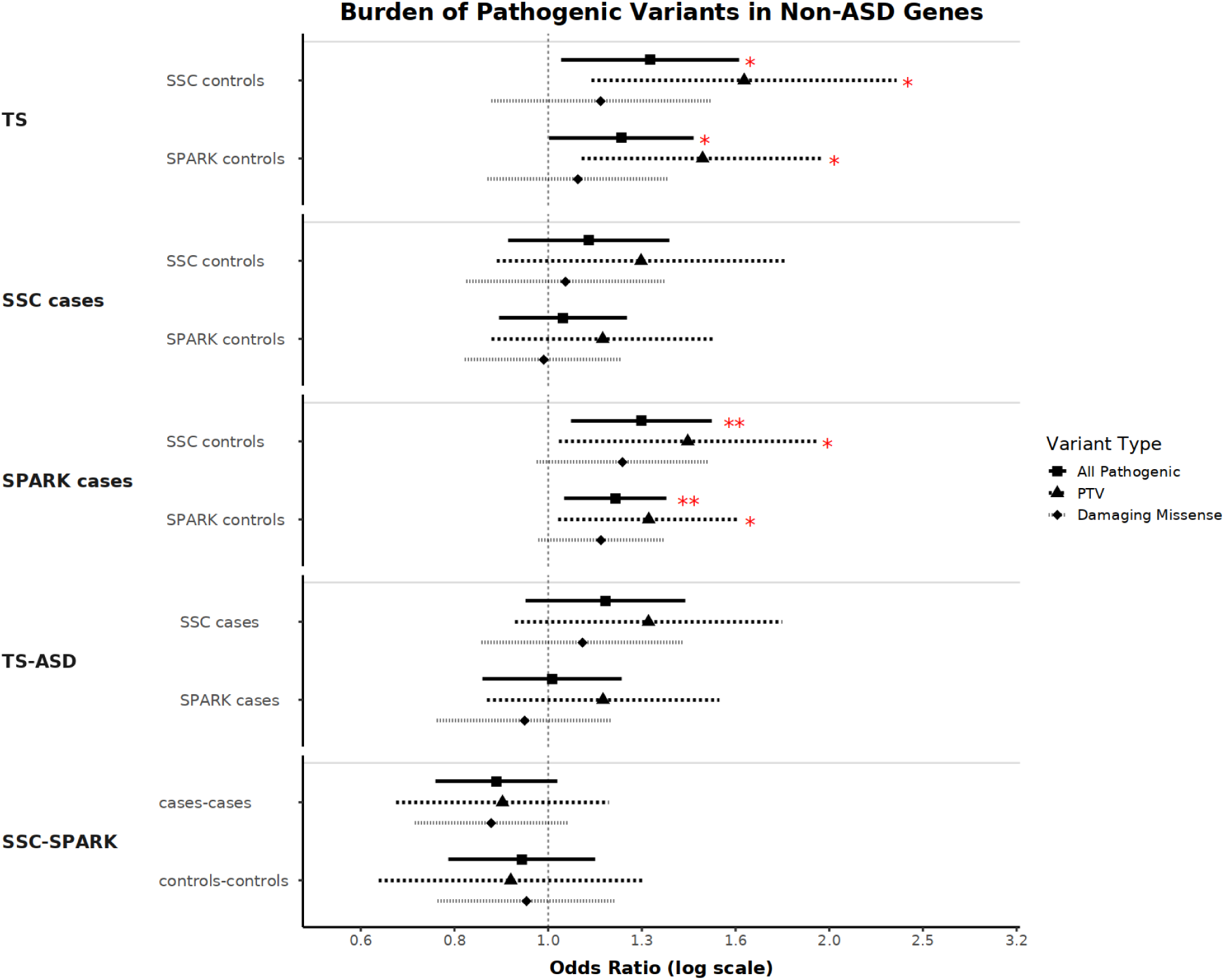
Forest plot of the burden analysis of DNMs between cohorts with the known ASD genes removed, split by variant annotations. All pathogenic, PTV + Dmis; PTV, protein-truncating variants; Dmis, missense variants with a REVEL score ≥ 0.644.

**Table 7.** Burden analysis of DNMs between cohorts with the known ASD genes removed, split by variant annotations. All pathogenic, PTV + Dmis; PTV, protein-truncating variants; Dmis, missense variants with a REVEL score ≥ 0.644.

| Comparison |  | All Pathogenic |  | PTV |  | Dmis |  |
| --- | --- | --- | --- | --- | --- | --- | --- |
| Sample 1 | Sample 2 | OR | Pvalue | OR | Pvalue | OR | Pvalue |
| TS | SSC Controls | 1.28 | 2.45E-02 | 1.62 | 1.17E-02 | 1.14 | 3.48E-01 |
|  | SPARK Controls | 1.20 | 4.78E-02 | 1.46 | 1.22E-02 | 1.08 | 5.21E-01 |
| SSC cases | SSC Controls | 1.10 | 3.22E-01 | 1.26 | 2.07E-01 | 1.04 | 7.33E-01 |
|  | SPARK Controls | 1.04 | 6.51E-01 | 1.14 | 3.37E-01 | 0.99 | 9.12E-01 |
| SPARK cases | SSC Controls | 1.26 | 9.85E-03 | 1.41 | 3.35E-02 | 1.20 | 8.96E-02 |
|  | SPARK Controls | 1.18 | 9.65E-03 | 1.28 | 2.95E-02 | 1.14 | 9.62E-02 |
| TS | SSC Cases | 1.15 | 1.60E-01 | 1.28 | 1.41E-01 | 1.09 | 5.06E-01 |
|  | SPARK Cases | 1.01 | 9.11E-01 | 1.14 | 3.56E-01 | 0.94 | 5.99E-01 |
| SSC-SPARK | Cases-Cases | 0.88 | 9.61E-02 | 0.89 | 4.00E-01 | 0.87 | 1.41E-01 |
|  | Controls-Controls | 0.94 | 4.81E-01 | 0.91 | 5.80E-01 | 0.95 | 6.31E-01 |

To explore the impact of removing DD risk genes, we repeated these analyses in all three cohorts after excluding these genes. Similar to the previous observation, the enrichment of pathogenic DNMs in the SSC ASD probands was eliminated, while we still observed a significant remaining burden of PTVs (OR = 1.41∼1.49, P-value = 1.13e-2∼2.31e-2) in the TS probands, though with smaller effect sizes compared to the exome-wide burden. Similarly, we observed significant burdens of both pathogenic DNMs and PTVs in the TS probands (OR = 1.21∼1.61, P-value = 6.49e-3∼4.10e-2), as well as in the SPARK ASD probands (OR = 1.19∼1.46, P-value = 4.64e-3∼1.77e-2), supporting a distinct TS genetic etiology from DD. (Figure 8 and Table 8) Notably, the significant depletion of DNMs in known ASD and DD genes from TS probands was absent in the remaining exome, providing secondary evidence to support a distinctive set of core TS genes. The analysis utilizing missense variants with other REVEL thresholds are included in Supplementary Figure 9-12.

**Figure 8.**
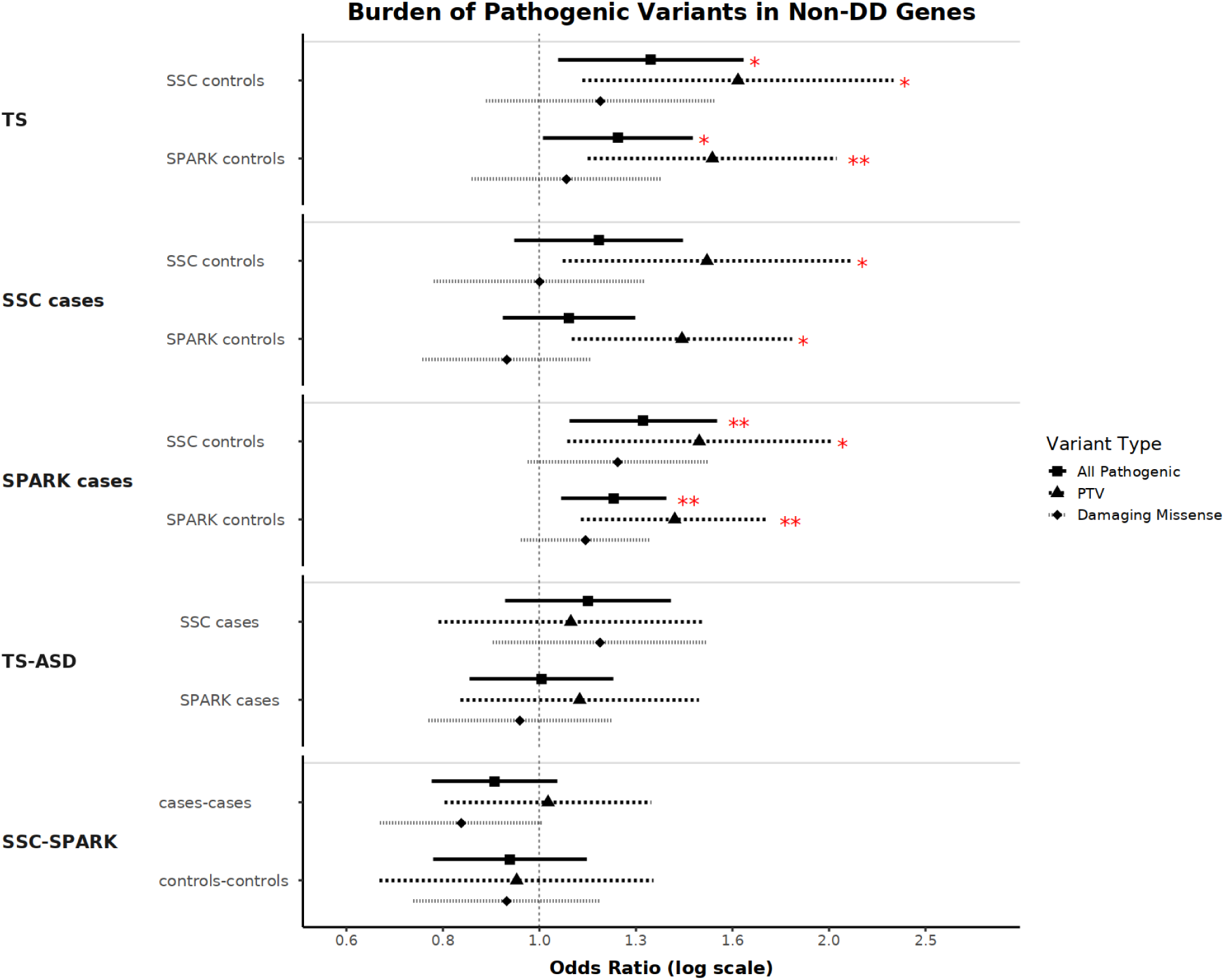
Forest plot of the burden analysis of DNMs between cohorts with the known DD genes removed, split by variant annotations. All pathogenic, PTV + Dmis; PTV, protein-truncating variants; Dmis, missense variants with a REVEL score ≥ 0.644.

**Table 8.** Burden analysis of DNMs between cohorts with the known DD genes removed, split by variant annotations. All pathogenic, PTV + Dmis; PTV, protein-truncating variants; Dmis, missense variants with a REVEL score ≥ 0.644.

| Comparison |  | All Pathogenic |  | PTV |  | Dmis |  |
| --- | --- | --- | --- | --- | --- | --- | --- |
| Sample 1 | Sample 2 | OR | Pvalue | OR | Pvalue | OR | Pvalue |
| TS | SSC Controls | 1.30 | 1.88E-02 | 1.61 | 1.25E-02 | 1.16 | 2.97E-01 |
|  | SPARK Controls | 1.21 | 4.10E-02 | 1.51 | 6.49E-03 | 1.07 | 5.75E-01 |
| SSC cases | SSC Controls | 1.15 | 1.67E-01 | 1.49 | 2.31E-02 | 1.00 | 9.98E-01 |
|  | SPARK Controls | 1.07 | 3.82E-01 | 1.41 | 1.13E-02 | 0.93 | 4.47E-01 |
| SPARK cases | SSC Controls | 1.28 | 6.03E-03 | 1.46 | 1.77E-02 | 1.21 | 8.80E-02 |
|  | SPARK Controls | 1.19 | 5.91E-03 | 1.38 | 4.64E-03 | 1.12 | 1.63E-01 |
| TS | SSC Cases | 1.12 | 2.51E-01 | 1.08 | 6.42E-01 | 1.16 | 2.68E-01 |
|  | SPARK Cases | 1.00 | 9.55E-01 | 1.10 | 5.07E-01 | 0.95 | 6.74E-01 |
| SSC-SPARK | Cases-Cases | 0.90 | 1.62E-01 | 1.02 | 8.70E-01 | 0.83 | 5.86E-02 |
|  | Controls-Controls | 0.93 | 4.50E-01 | 0.95 | 7.45E-01 | 0.92 | 4.89E-01 |

### Integration of both DNMs and rare inherited variants identifies three TS risk genes

The relative risk of variants has been previously reported to be differentially dependent on the mode of inheritance, variant classes, and evolutionary constraint.(42) Using a previously published Bayesian analytic framework, TADA(41), we combined these variables into one unified metric to maximize our power to discover individual risk genes in TS. We calculated a Bayes factor (BF) for each autosomal protein-coding gene by incorporating both DNMs and rare inherited variants, as well as accounting for mutation rates and relative risk priors adopted from the previous study(42).

This unified framework, applied to the TS probands and unaffected siblings from the two ASD cohorts, resulted in the identification of three TS risk genes reaching the threshold of FDR ≤ 0.05 (*PPP5C*, *EXOC1* and *GXYLT1*). We further tabulated the number of rare PTV and Dmis variants in both the *de novo* and the inherited categories in each of these three risk genes. None of the three genes had an excessive amount of PTV or Dmis variants.

## Discussion

This study represents one of the largest whole-exome sequencing efforts aimed at investigating the genetic architecture of Tourette syndrome (TS) to date, integrating data from 4,430 individuals from TS families (1,466 trios), 9,449 individuals from the Simons Simplex Cohort (SSC) (2,483 quads) and 16,924 individuals from the Simons Powering Autism Research (SPARK) cohort (4,231 quads). By leveraging an analytical framework that incorporates assessment of both *de novo* and rare inherited variants, we identified significant contributions of rare coding variants to TS risk, particularly protein-truncating variants (PTVs) and damaging missense variants (Dmis). These findings provide new insights into the genetic etiology of TS and highlight both shared and distinct features compared to ASD and DD.

### Rare and De Novo Variants in TS and ASD

Our results demonstrate a significant exome-wide enrichment of pathogenic DNMs in TS probands, with PTVs being the primary driver of this burden. This observation aligns with previous studies in neurodevelopmental and neuropsychiatric disorders, where PTVs have been shown to confer substantial risk due to their disruptive effects on gene function.(21,33,49–52) The enrichment of DNMs in TS was particularly pronounced in loss-of-function (LoF)-intolerant genes (pLI ≥ 0.9 and LOEUF < 0.6), suggesting that these mutations disproportionately affect genes critical for human development and neuronal function. This pattern mirrors findings in ASD, where DNMs in LoF-intolerant genes have been consistently associated with increased risk.(42,53) However, unlike ASD, we observed only a trend for enrichment of Dmis variants in TS probands that was not statistically significant, indicating that PTVs may play a more dominant role in TS genetic architecture than do Dmis variants and that additional samples might be required to further elucidate the specific contribution of Dmis variants in TS.

### Shared and Distinct Genetic Architectures

TS is highly polygenic, and previous studies indicate that both common and rare variants contribute to TS risk.(7,12,21–23,33,54) Previous exome sequencing studies have also demonstrated that rare coding variants, particularly PTVs, play a significant role in ASD risk.(27,55) Although there is some evidence of clinical overlap between TS and ASD, as well as shared polygenic risk, a key finding of our study is the fact that we found limited convergence between TS and ASD in terms of risk from rare genetic variants. (8,34,38,56,57) Consistent with prior work, we first observed significant enrichment of DNMs in ASD probands within known ASD and DD risk genes—gene sets largely defined from cohorts such as SSC and SPARK—thereby validating the quality and robustness of our variant-calling and analytical pipeline. In contrast, TS probands showed no enrichment in these previously implicated ASD/DD genes, while retaining significant DNM enrichment elsewhere in the genome. Despite substantial clinical overlap, previous studies have estimated that the genetic correlation between TS and ASD is between 0.1 and 0.2, compared to 0.4 for TS and OCD, another childhood-onset neurodevelopmental disorder with substantial TS overlapping features.(12,58) These findings, taken together, suggest that the genetic underpinnings of TS are distinct from those of ASD and DD, with limited overlap between the two disorders, at least with regard to the contributions of rare variants, implying that TS risk genes might lie in non-ASD/DD loci. This divergence highlights the unique genetic contributions to TS and the distinct molecular pathways and neural circuits that may be involved in this developmental neuropsychiatric disorder.

Interestingly, the SPARK ASD cohort exhibited a different genetic architecture compared to the SSC ASD cohort, with pathogenic DNMs and PTVs continuing to contribute significantly to ASD risk even after the exclusion of ASD risk genes. This heterogeneity may reflect the broader phenotypic spectrum captured in the SPARK cohort, which includes individuals with comorbid neuropsychiatric conditions, consistent with previous literature.(59,60) These findings emphasize the need for careful consideration of cohort composition and phenotypic heterogeneity in genetic studies of neuropsychiatric disorders, as well as the importance of understanding their shared and distinct genetic architectures.

### Identification of Novel TS Risk Genes

Using a Bayesian statistical framework (TADA), we integrated the *de novo* and rare inherited variants that met our threshold for statistical significance, and identified three novel TS risk genes: *PPP5C*, *EXOC1*, and *GXYLT1.* These genes were associated with TS at a false discovery rate (FDR) ≤ 0.05. The *PPP5C* gene encodes a protein from the protein phosphatase catalytic subunit family and participates in pathways regulated by reversible phosphorylation, essential to cell growth and differentiation. Its expression is high in the mammalian brain.(61–65) Previous literature has associated *de novo* mutations in this gene with microcephaly, seizures, and developmental delay.(66) *PPP5C* has also been identified in ASD studies examining rare *de novo* transmitted variants and in GWAS of pain-related phenotypes.(67,68) The *EXOC1* gene encodes a highly conserved protein that is part of the exocyst complex involved in the targeting of exocytic vesicles to the plasma membrane. This gene has been linked to psychosis liability, and its interaction with *DISC1* has been implicated in major mental illnesses, such as schizophrenia.(69–71) Other subunits of the exocyst complex, in the same family as *EXOC1*, have also been associated with a range of neurodevelopmental disorders.(72–76) The last identified TS risk gene, *GXYLT1*, encodes a xylosyltransferase. This gene has been linked to completed (fatal) suicide and major depression disorder in previous GWAS and to post-traumatic stress disorder in gene expression analysis.(77–79) While the direct functional relevance of these genes to TS remains to be fully elucidated, many of the associated neuropsychiatric conditions found in the literature are genetically correlated with TS. Hence, their identification represents a significant step forward in understanding the genetic basis of TS. Future studies with larger sample sizes and functional validation will be necessary to confirm their roles in TS pathogenesis and to explore potential biological pathways and functional consequences involved, shedding new light into the neurobiological mechanisms underlying TS.

### Limitations and Future Directions

While this study represents a significant advance in our understanding of the genetic basis of TS, several limitations should be acknowledged. First, the sample size of the TS cohort, though one of the largest to date, remains smaller than those of the SSC and SPARK cohorts, which may limit our power to detect additional risk genes and fully elucidate the genetic contributions from specific variant groups. One such example is the Dmis variants. The wide confidence intervals (CIs) and standard errors (SEs) of the Dmis variants observed in this study, compared to those observed for PTVs variants, indicate that a larger sample size is required to reach statistical significance and/or confidence in these findings, given that the ORs were above 1 for various scopes tested in our TS burden analysis, suggesting a potentially noticeable contribution of Dmis variants to TS risk. Second, phenotypic heterogeneity within the data, such as SPARK, and the inclusion of individuals with comorbid conditions, especially in the control samples, may introduce noise into our analyses and reduce our power to detect causal associations. On the other hand, the use of “super-controls”, i.e., individuals without any comorbid conditions, could also bias results, as such individuals are unrepresentative of the general population and may exaggerate differences between cases and controls. Future studies with larger and diagnostically more homogeneous cohorts will be essential to refine these findings and identify additional TS risk genes.

Additionally, the functional annotation of rare variants, particularly missense variants, remains challenging. While we used REVEL scores to classify Dmis variants, the biological impact of these variants may vary depending on their specific location and effects on protein function. Integrating additional functional annotations, such as gene expression data and protein-protein interaction networks, may improve our ability to prioritize candidate genes and variants.

## Conclusion

In summary, this study provides new insights into the genetic architecture of TS, highlighting the significant contribution of rare *de novo* coding variants, particularly PTVs, to TS risk. We identified three novel TS risk genes and demonstrated that the genetic underpinnings of TS, at least with respect to the contribution from rare variants identified by whole exome sequencing, are distinct from those of ASD and DD. These findings underscore the importance of studying TS as a unique neuropsychiatric disorder and provide a foundation for future research into its neurobiological mechanisms. As sample sizes continue to grow and analytical methods improve, we anticipate further discoveries that will deepen our understanding of TS and inform the development of targeted therapies and pharmacological solutions.

## Methods

### Sample collection

A total of 1500 proband-parent trios were recruited from 14 sites across the United States, Canada, the United Kingdom, the Netherlands, Israel, Costa Rica, and Colombia. Probands met inclusion criteria defined by the Tourette Syndrome Classification Study Group (TSCSG), requiring a diagnosis of definite TS,corresponding to DSM-IV-TR criteria and confirmation of tics by an experienced clinician. Exclusion criteria included a history of intellectual disability (ID), ASD, or tardive tourettism. Blood samples or cell lines were collected from probands and their parents for genomic DNA extraction.

ASD samples and controls were collected from two cohorts: 1). the Simons Foundation Autism Research Initiative (SFARI) Simons Simplex Collection (SSC); 2). the Simons Foundation Powering Autism Research for Knowledge (SPARK). The SSC cohort contains 9,949 samples from over 2,000 families and is clinically evaluated to contain only probands with moderate to severe symptoms, as well as their healthy siblings and parents.(80) The SPARK cohort was recruited through online platforms and the iWES v1 release contains 70,489 samples from over 30,000 families.(81) To maintain the same study design, we pre-selected the families with at least one proband, one healthy sibling, and both parents available, which resulted in a total number of 16,924 samples.

All cohorts had accompanying whole-exome sequencing data, sequenced from multiple platforms and with different targets. To homogenize the data and minimize the impact of technical differences, we jointly processed and genotyped all three cohorts following the GATK best practices.(82) Briefly, we obtained the target interval lists from the three cohorts and extracted only the intersecting regions. To enhance the calling accuracy of the slice-site variant, we added 100 bp paddings around each interval. Within the padded regions, we then generated gVCF files from raw cram files using the GATK HaplotypeCaller in gVCF mode, followed by a consolidation step with GenomicsDBImport. The resulting data were then jointly genotyped for high-confidence alleles using GenotypeGVCFs, which were then calibrated and filtered using VariantRecalibrator and ApplyVQSR.

### Variant- and sample-level filtering

To minimize the impact of technical factors and include only high-quality variants, we applied stringent QC filters to the jointly called dataset. The VCFtools v0.1.16(83) was used to process the VCF files and produce the final working dataset. In brief, the variants were assessed in three QC steps. First, only variants that satisfied the following criteria were retained: 1). bi-allelic variants; 2). variant type is SNP; 3). variants with alternative AC ≥ 1; 4). variants annotated with the PASS flag by the VQSR model; 5). genotypes with GQ ≥ 21. This minimal QC step focused on cleaning up variant types irrelevant to this study and filtered out low-quality variants with pre-calculated annotations. In the second step, we looked at the distribution of the variant missing rate across the three cohorts, and generated a combined list of SNPs with a missing rate exceeding 5% in each cohort, which ensured the heterogeneity in the variant missingness due to sequencing batches was minimized. Lastly, to further eliminate potentially low-quality variants, we set the following genotypes to missing: 1). DP < 10; 2). AB_het < 0.2, and then filtered out any variants with new missing rates > 10% and HWE p-values < 1×10^−12^ in the resulting data.

Similar to variant-level filtering, sample-level filtering was also performed to remove samples with overall low sequencing quality. After variants with high missing rates were filtered out, samples that remained to have missing rates > 10% were left out. The identity-by-descent (IBD) analysis was performed to check the relatedness within and across the three cohorts. The analysis was performed with the tool Kinship-based INference for Genome-wide association studies (KING) v.2.3.0, and one sample from each pair of duplicates or unexpected first-degree relatives was excluded based on the missing rate. The pedigree structures of the three cohorts recorded in the manifest files were also attested against the sample relatedness inferred from the genetic distance. Sex check analysis was performed using PLink v1.9(84), and all samples with discordant sex predictions were examined based on pedigree structures. Subsequently, the sex records were corrected, and the samples were retained only when the pedigree structures of the affected families were not violated. Additionally, we estimated the concordance rate between the array genotype data and the WES data from the TS cohort. A total of ten samples were removed due to low concordance rates (< 95%).

### Variant calling and quality control of inherited and DNMs

To select candidates for *de novo* variant calling, we obtained rare variants from the total set of high-quality variants that passed the QC filters mentioned in the previous section. We treated variants in two different ways. On the one hand, for the variants present in the gnomAD v2.1.1 liftover dataset (GRCh38), we defined rare variants to have minor allele counts (MACs) ≤ 20 within the non-Neuro subset of the samples (N=104,064), which was equivalent to a threshold of MAF ≤ 0.01%. On the other hand, if the variants were unique to our dataset, we required rare variants to have MACs ≤ 5 (equivalent to MAF ≤ 0.0024%). Additionally, to further ensure the sequencing quality of the candidates, we evaluated the allele balance of the homozygous (AB_hom) and the heterozygous (AB_het) genotypes; only homozygous genotypes with AB_hom ≤ 0.05 and heterozygous genotypes with AB_het between 0.3 and 0.7 were kept.

To call DNMs from the curated candidates, we used the tool TrioDeNovo v0.06(85), a Bayesian framework specifically designed to call DNMs in trios with flexible prior assignments based on genomic regions. One inconvenient feature of this tool was that it required all input families to be strictly trios. Therefore, out of 7,641 unique families from our joint-calling dataset, we created pseudo-families by duplicating parents for the families with more than one child, which resulted in a total of 14,046 trios. We set the parameters minDQ and minDP to be 7 and 5, respectively. The former metric was estimated by TrioDeNovo to assess the minimum quality scores for the called *de novo* variant to be included, while the latter required the read depth of a given *de novo* variant to be over 10 in all three individuals (father, mother, and child) from the respective trio.

### Variant annotation

Putative DNMs were annotated using the Variant Effect Predictor (VEP) version 106. Variants were classified into four categories: modifiers, low impact, moderate impact, and high impact. Variants with impact levels below moderate were removed from further analysis, except for synonymous variants, which were used for baseline mutation rate estimation in both cases and controls. High-impact variants included protein-truncating variants, as well as stop-loss, splice-acceptor, and splice-donor variants. Moderate variants included missense, in-frame insertion, in-frame deletion, and protein-altering variants. To ensure consistent annotation across the entire dataset, all identified DNMs were processed through this pipeline.

For further characterization of missense mutations, following the guide of a recent study on missense variant pathogenicity classification for clinical interpretation, we further annotated missense variants with REVEL scores.(44) Predicted pathogenicity was assessed using three recommended thresholds: ≥0.932, ≥0.773, and ≥0.644, with the latter including all variants from the former. The last category included all missense variants from the other two categories and was defined as the damaging missense variants (Dmis) category for further analysis. To note, one variant that truncates the last exons of *FMO2* was found to be the major allele in the cohort and was predicted to be “likely benign” by the Clinvar database, suggesting a limited functional impact. Hence, we removed it from further analysis. All other genes with at least one pathogenic variant, defined as either PTV or Dmis variant, were kept for later analyses.

### TADA Bayesian framework to identify risk genes

To identify individual risk genes for TS, we employed the Transmission and De Novo Association (TADA) model, a Bayesian statistical framework.(86) TADA is designed to measure disease association at the gene level by integrating data from different types of genetic variants, including inherited rare variants and DNMs. For each autosomal protein-coding gene, the model calculates a Bayes Factor (BF), incorporating the observed counts of specific variant events (in our case, de novo mutations and rare inherited variants), gene-specific mutation rates, the number of samples analyzed, and prior probabilities regarding the risk of variants in the corresponding gene.

We leveraged the key strength of the TADA framework and aggregated BFs from distinct variant categories (namely, *de novo* PTVs, *de novo* Dmis variants, inherited rare PTVs, and inherited rare Dmis) for the same gene, which was subsequently used to estimate a False Discovery Rate (FDR). A statistical significance threshold of FDR≤0.05 was used to identify candidate risk genes.

### Burden test

All statistical analyses were performed using R statistical software (version 4.3.0).(87) Comparisons of DNM rates between TS probands and unaffected controls (from SSC and SPARK cohorts) were conducted using two-sample Poisson rate tests, as well as Wilcoxon rank sum tests. These analyses were applied exome-wide as well as to specific functional categories of DNMs, namely PTVs and Dmis variants, as defined in the “Variant annotation” section.

### Gene-set burden analysis

For gene-set burden analyses, assessing the burden of DNMs within specific gene categories like LoF-intolerant genes, or previously established lists of ASD and DD risk genes, we compared the proportion of DNMs occurring within a given gene set in TS probands against the proportion in unaffected controls from SSC and SPARK cohorts. We used, separately, two established metrics to define LoF-constrained genes: 1). pLI metric,(88) 2). LOEUF score.(43) The recommended threshold of pLI ≥ 0.9 was used to define LoF-intolerant genes. The LOUEF score had two versions based on the release (v2.1.1 and v4.0) of the underlying gnomAD data from which the score was calculated. As the former version was released earlier and thus established, the latter had an almost six-fold increase in sample size and used high-coverage bases to generate the LOUEF score. Hence, we included only the newer version in our analysis, using the recommended thresholds of LOEUF ≤ 0.6. To test the specificity of the association, we also removed ASD and DD gene risk genes from the total test set and assessed the strength of the association using the variants in the remaining genome. The P-values presented in the Results section correspond to these comparative analyses.

## Acknowledgement

We are deeply grateful to the individuals with Tourette syndrome (TS) and autism spectrum disorder (ASD), their families, and the volunteers who made this study possible. This work was supported by the National Institutes of Health (NIH) grant R01 NS102371 (to Drs. Scharf, Mathews, and Ophoff). TS family recruitment was conducted through the Tourette Syndrome Association International Consortium for Genetics (TSAICG), and ASD family recruitment was performed by the Simons Foundation. We thank the many investigators, clinicians, and technical staff who contributed to participant ascertainment, diagnostic evaluation, biospecimen processing, sequencing, and data curation. The funding organizations had no role in the design or conduct of the study; in the collection, management, analysis, or interpretation of the data; or in the preparation, review, or approval of the manuscript.

---

**Supplementary Figure 1.**
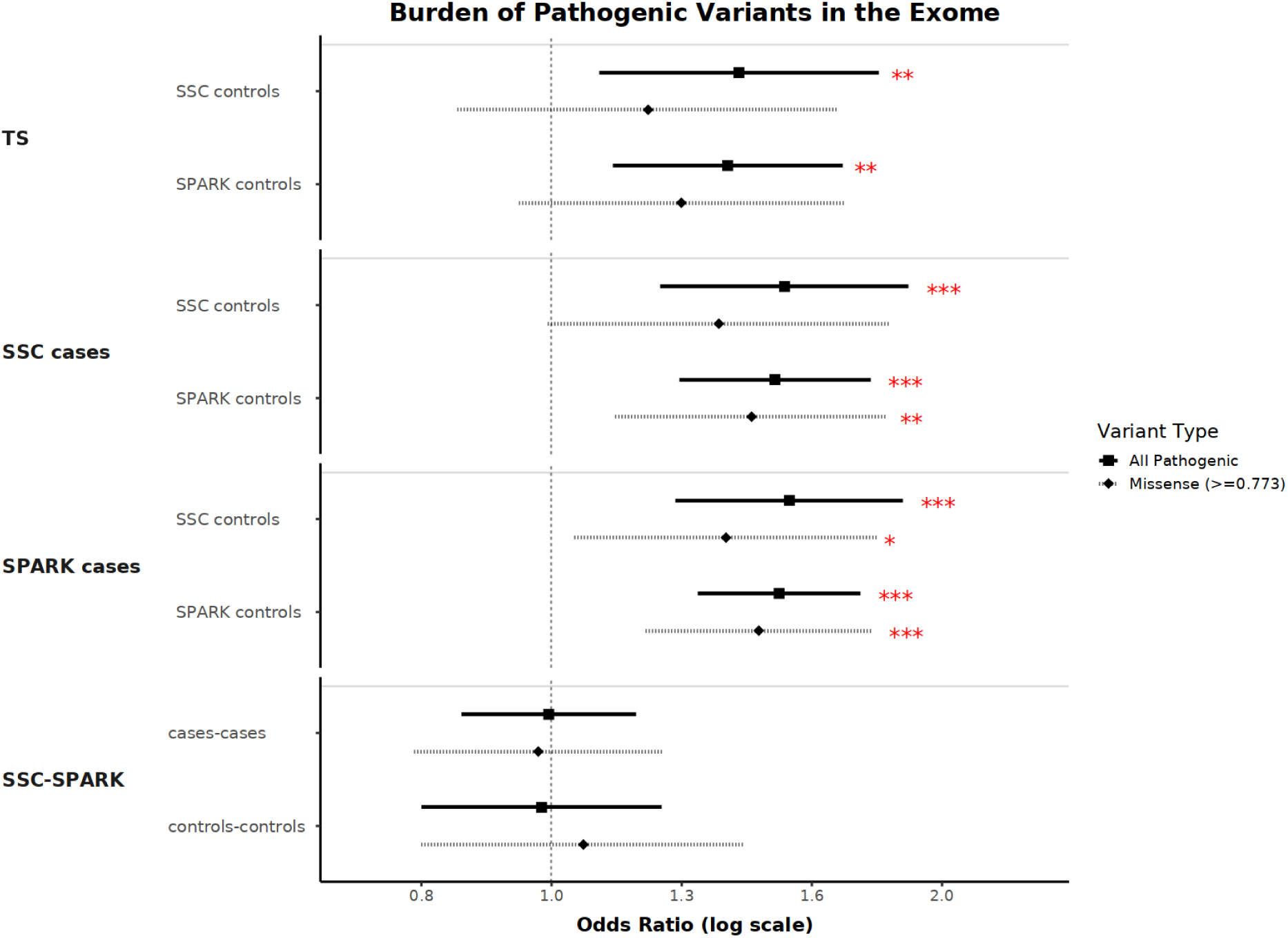
Forest plot of the burden analysis of DNMs between cohorts within the context of the exome, split by variant annotations. All pathogenic, PTV + Dmis; Dmis, missense variants with a REVEL score ≥ 0.773.

**Supplementary Figure 2.**
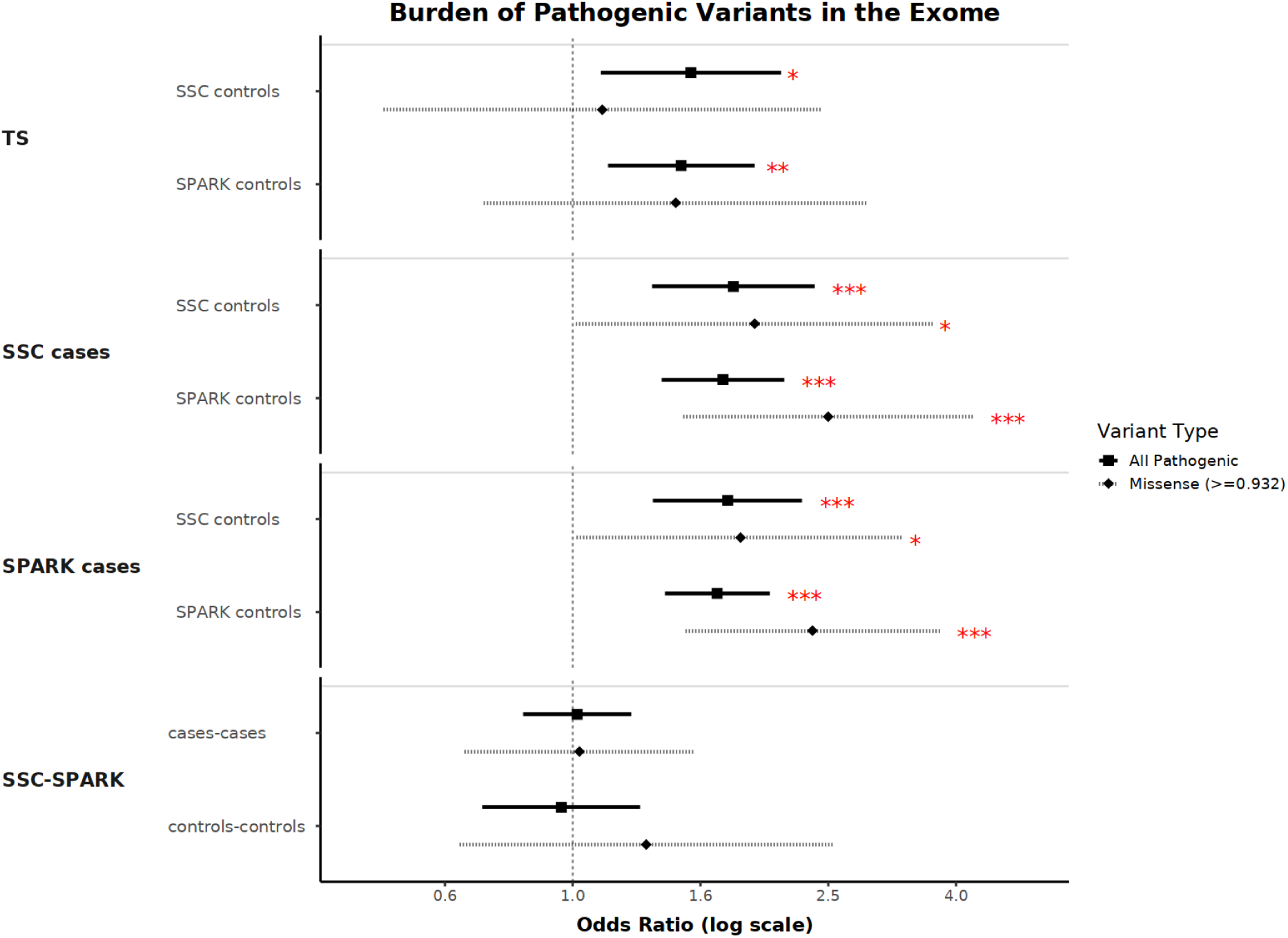
Forest plot of the burden analysis of DNMs between cohorts within the context of the exome, split by variant annotations. All pathogenic, PTV + Dmis; Dmis, missense variants with a REVEL score ≥ 0.932.

**Supplementary Figure 3.**
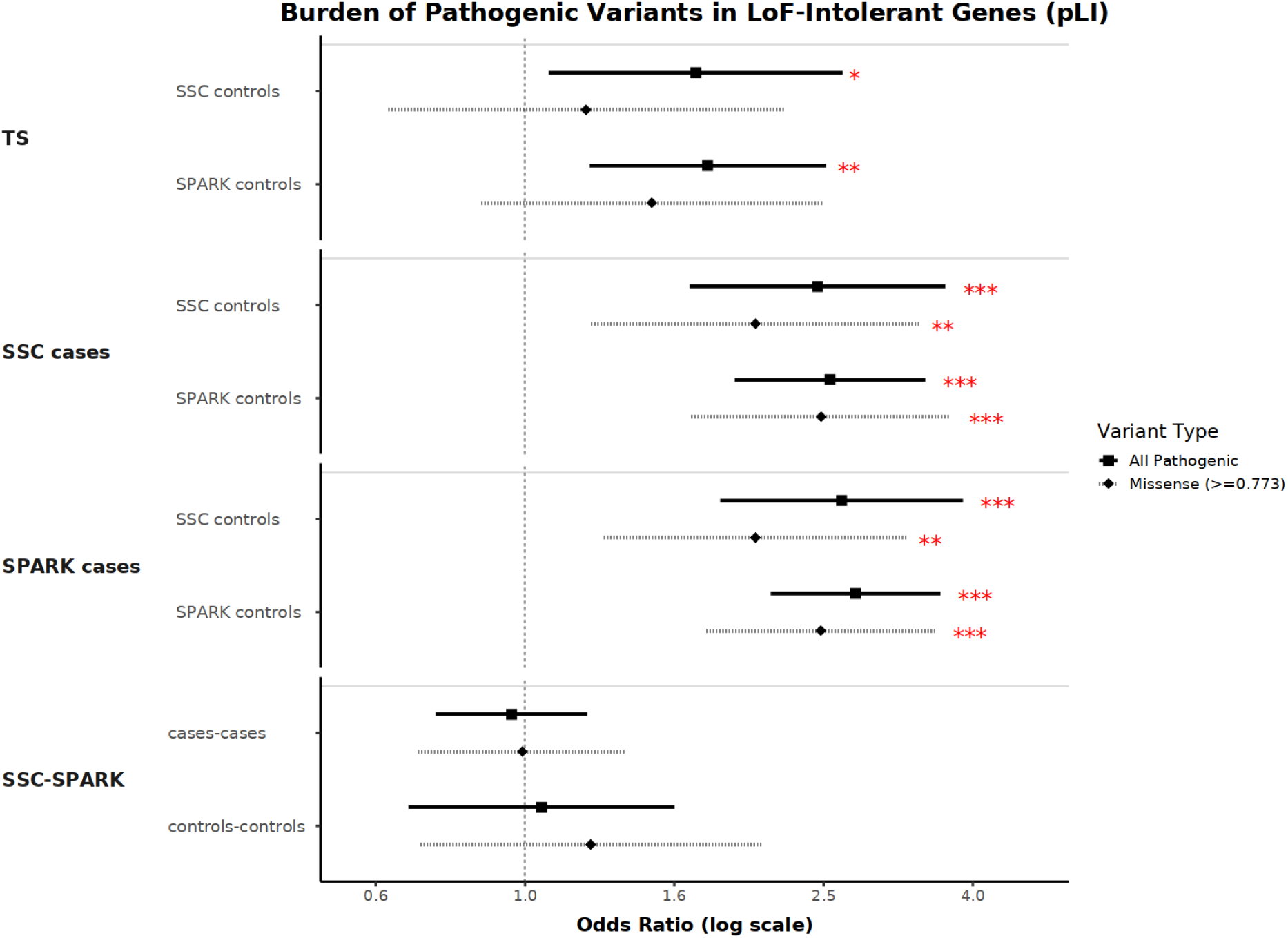
Forest plot of the burden analysis of DNMs between cohorts within the context of the LoF-intolerant genes (defined by pLI score), split by variant annotations. All pathogenic, PTV + Dmis; Dmis, missense variants with a REVEL score ≥ 0.773.

**Supplementary Figure 4.**
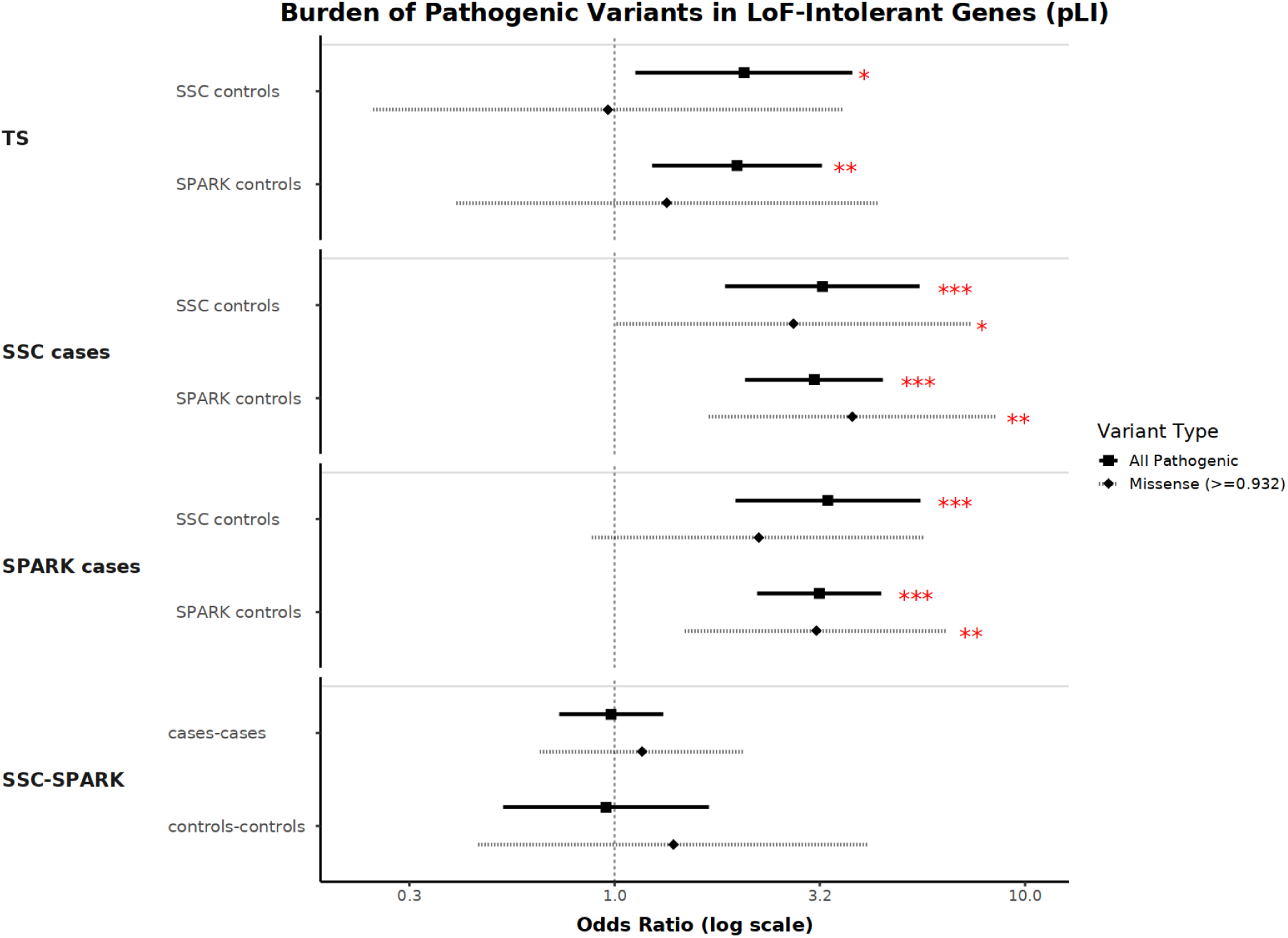
Forest plot of the burden analysis of DNMs between cohorts within the context of the LoF-intolerant genes (defined by pLI score), split by variant annotations. All pathogenic, PTV + Dmis; Dmis, missense variants with a REVEL score ≥ 0.932.

**Supplementary Figure 5.**
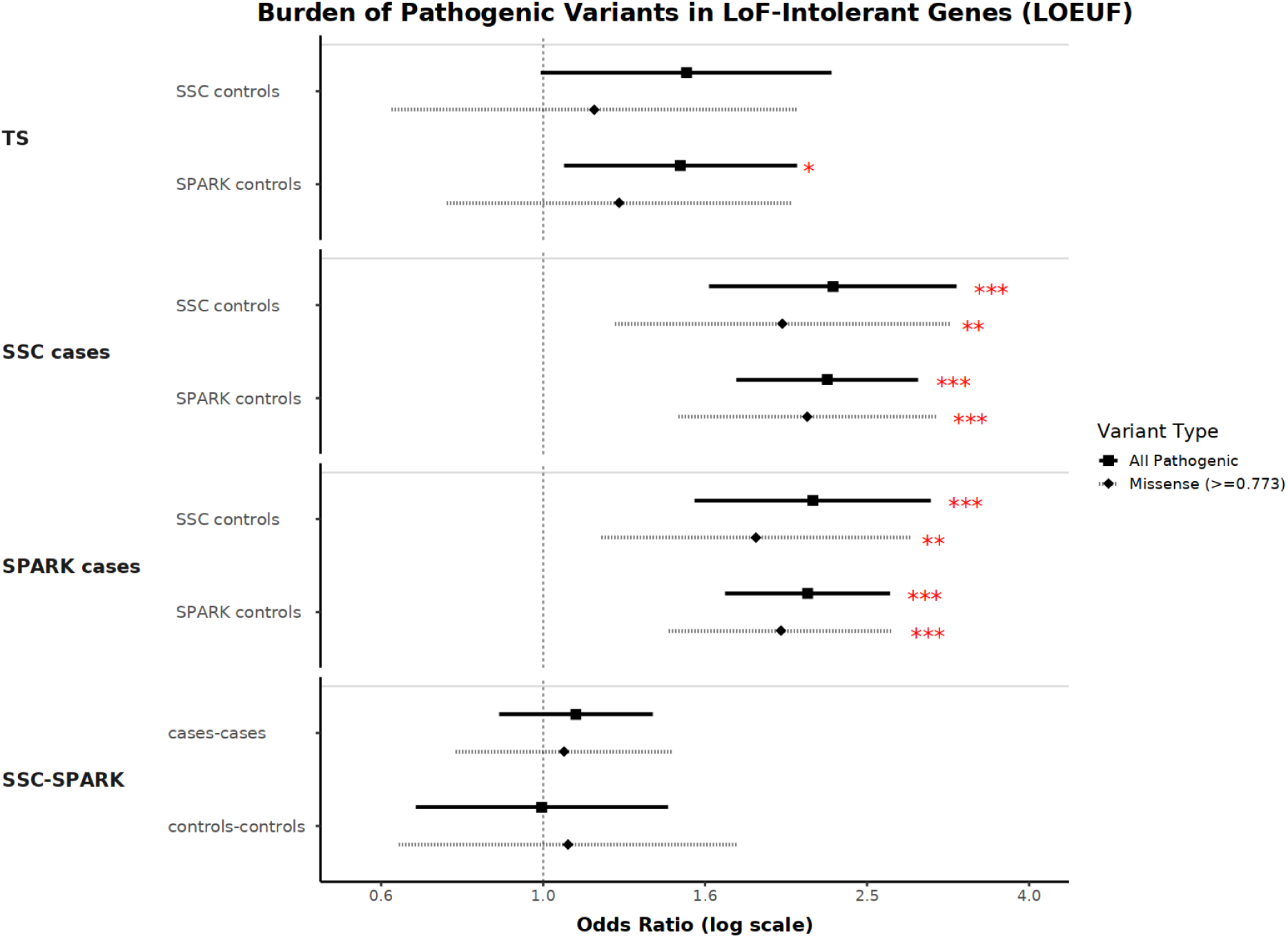
Forest plot of the burden analysis of DNMs between cohorts within the context of the LoF-intolerant genes (defined by LOEUF metric), split by variant annotations. All pathogenic, PTV + Dmis; Dmis, missense variants with a REVEL score ≥ 0.773.

**Supplementary Figure 6.**
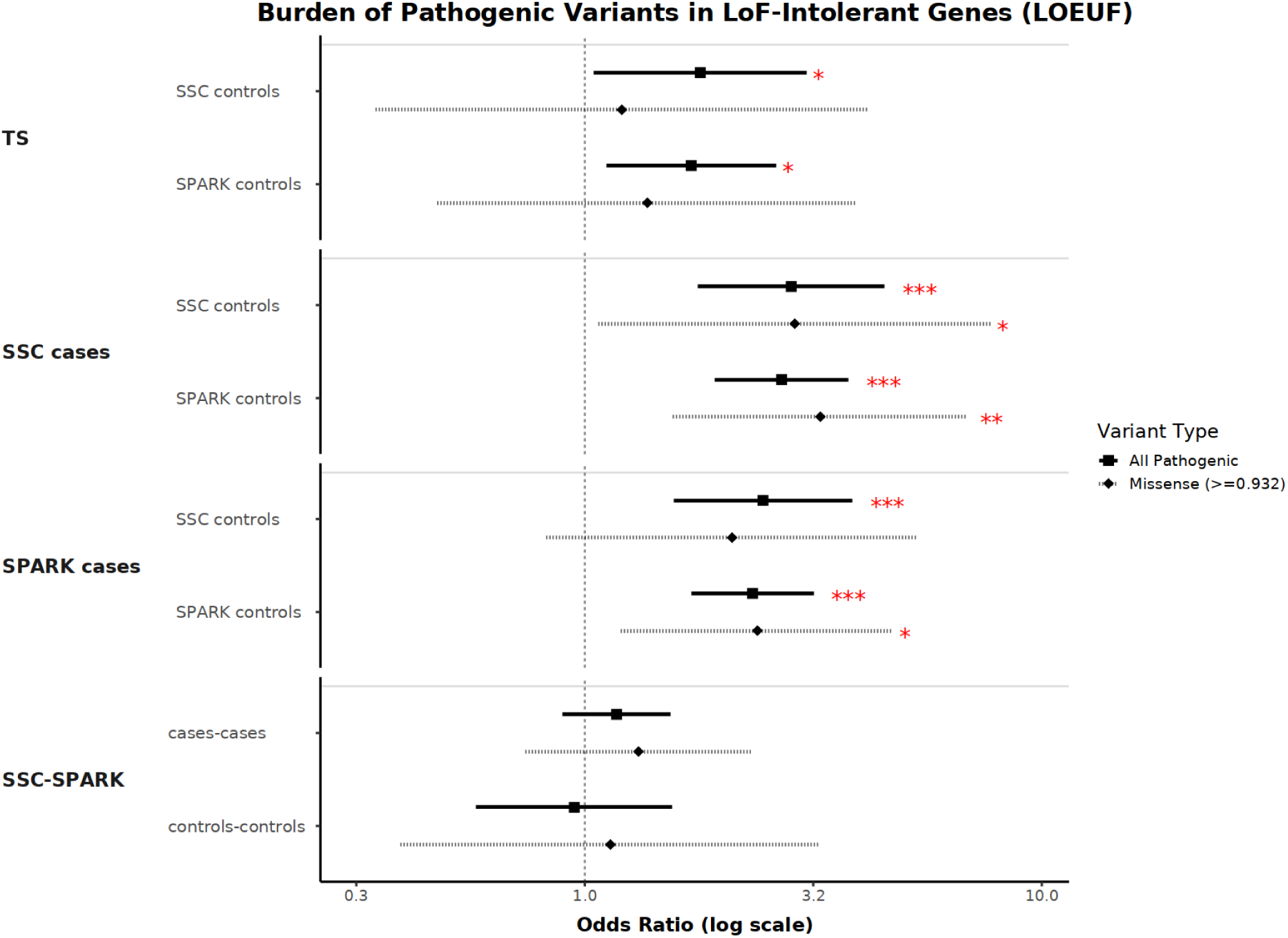
Forest plot of the burden analysis of DNMs between cohorts within the context of the LoF-intolerant genes (defined by LOEUF metric), split by variant annotations. All pathogenic, PTV + Dmis; Dmis, missense variants with a REVEL score ≥ 0.932.

**Supplementary Figure 7.**
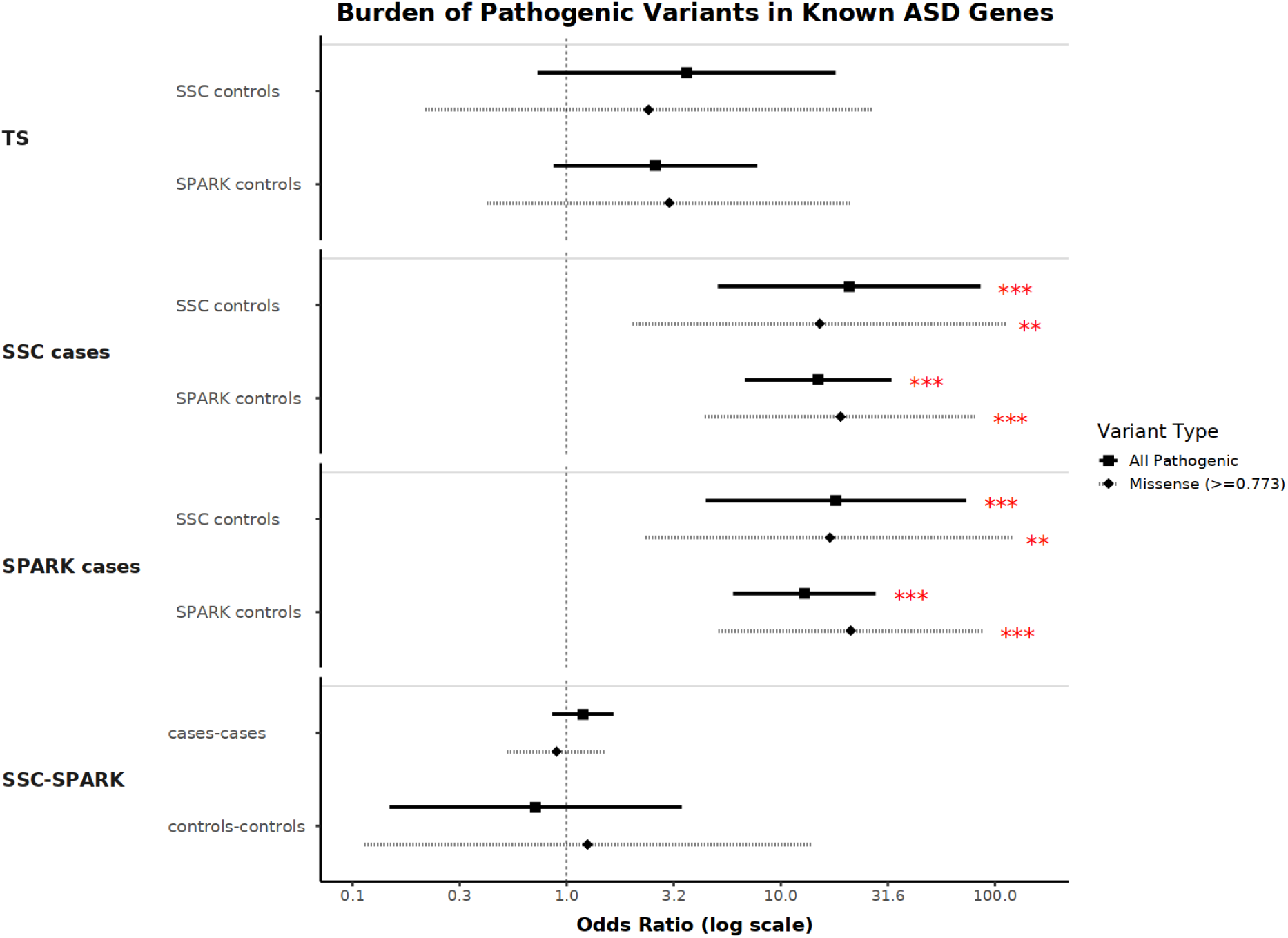
Forest plot of the burden analysis of DNMs between cohorts within the context of the known ASD genes, split by variant annotations. All pathogenic, PTV + Dmis; Dmis, missense variants with a REVEL score ≥ 0.773.

**Supplementary Figure 8.**
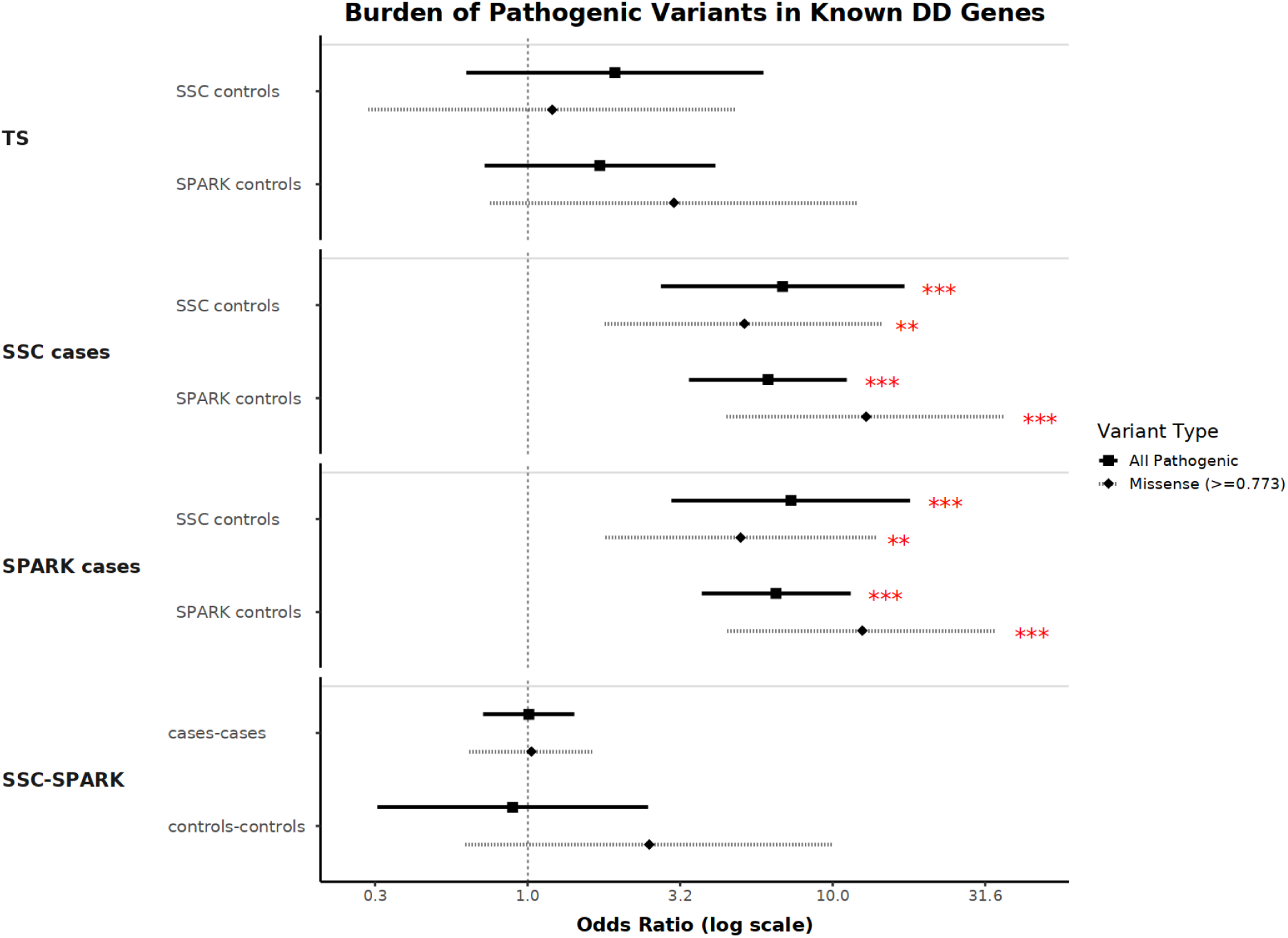
Forest plot of the burden analysis of DNMs between cohorts within the context of the known DD genes, split by variant annotations. All pathogenic, PTV + Dmis; Dmis, missense variants with a REVEL score ≥ 0.773.

**Supplementary Figure 9.**
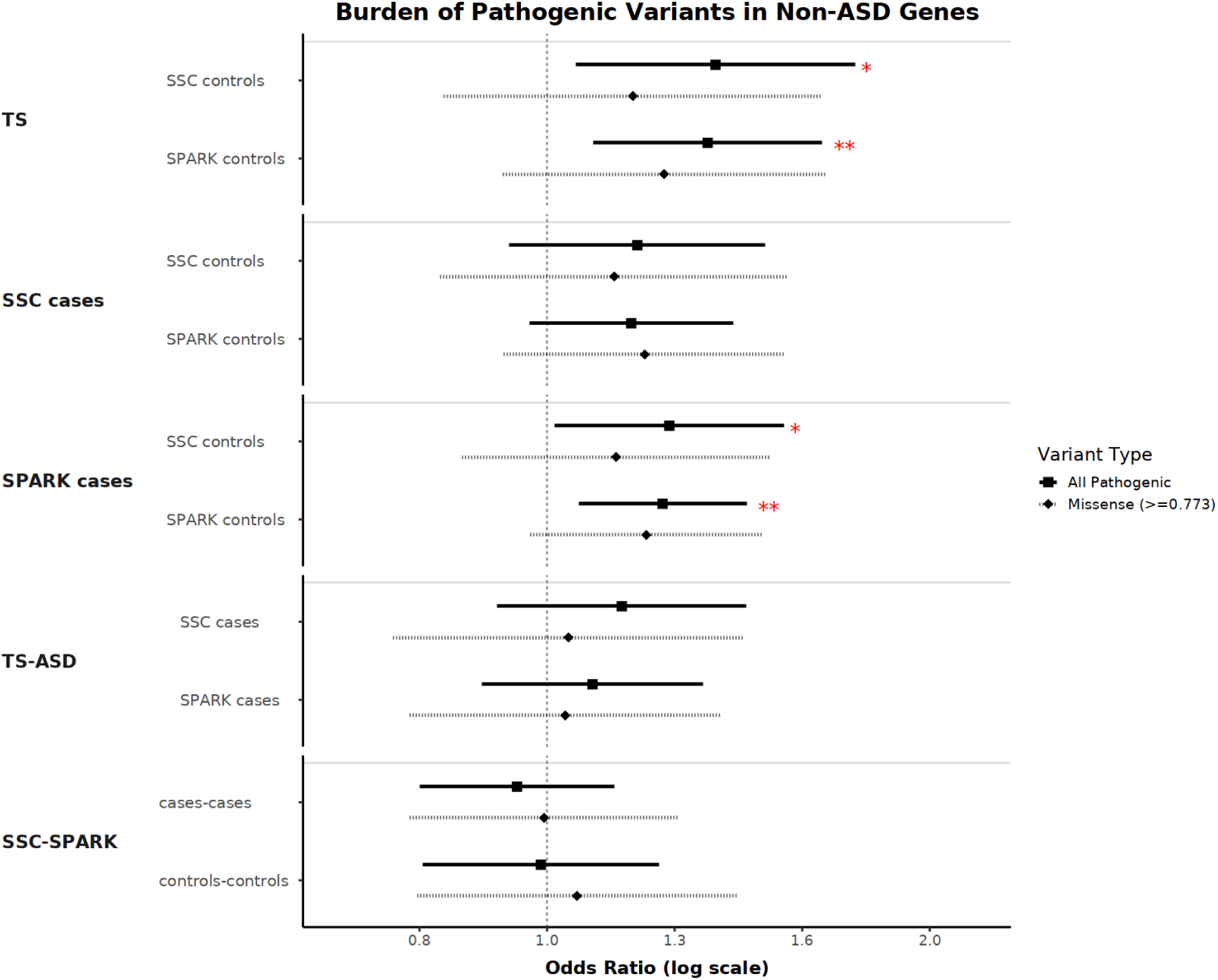
Forest plot of the burden analysis of DNMs between cohorts with the known ASD genes removed, split by variant annotations. All pathogenic, PTV + Dmis; Dmis, missense variants with a REVEL score ≥ 0.773.

**Supplementary Figure 10.**
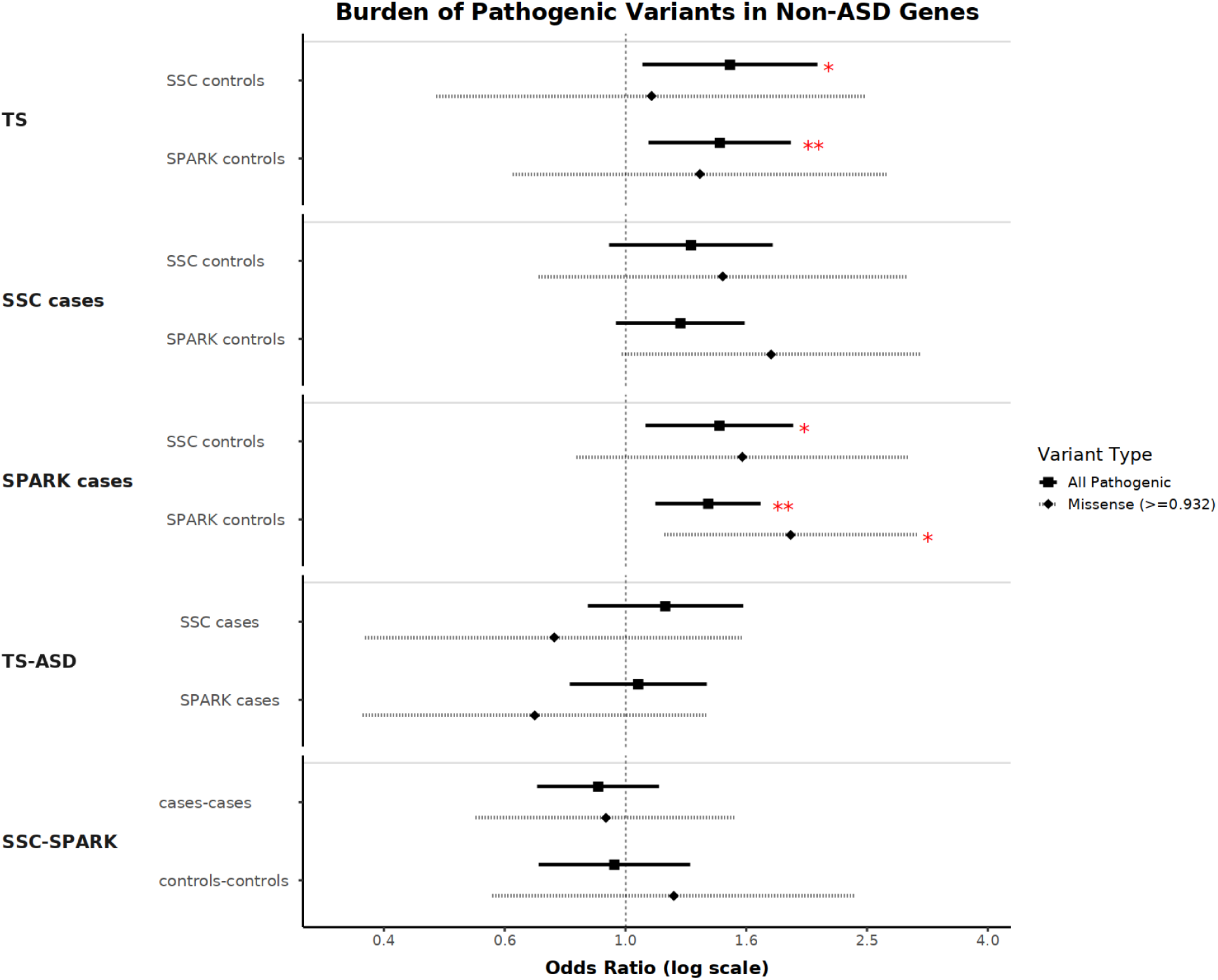
Forest plot of the burden analysis of DNMs between cohorts with the known ASD genes removed, split by variant annotations. All pathogenic, PTV + Dmis; Dmis, missense variants with a REVEL score ≥ 0.932.

**Supplementary Figure 11.**
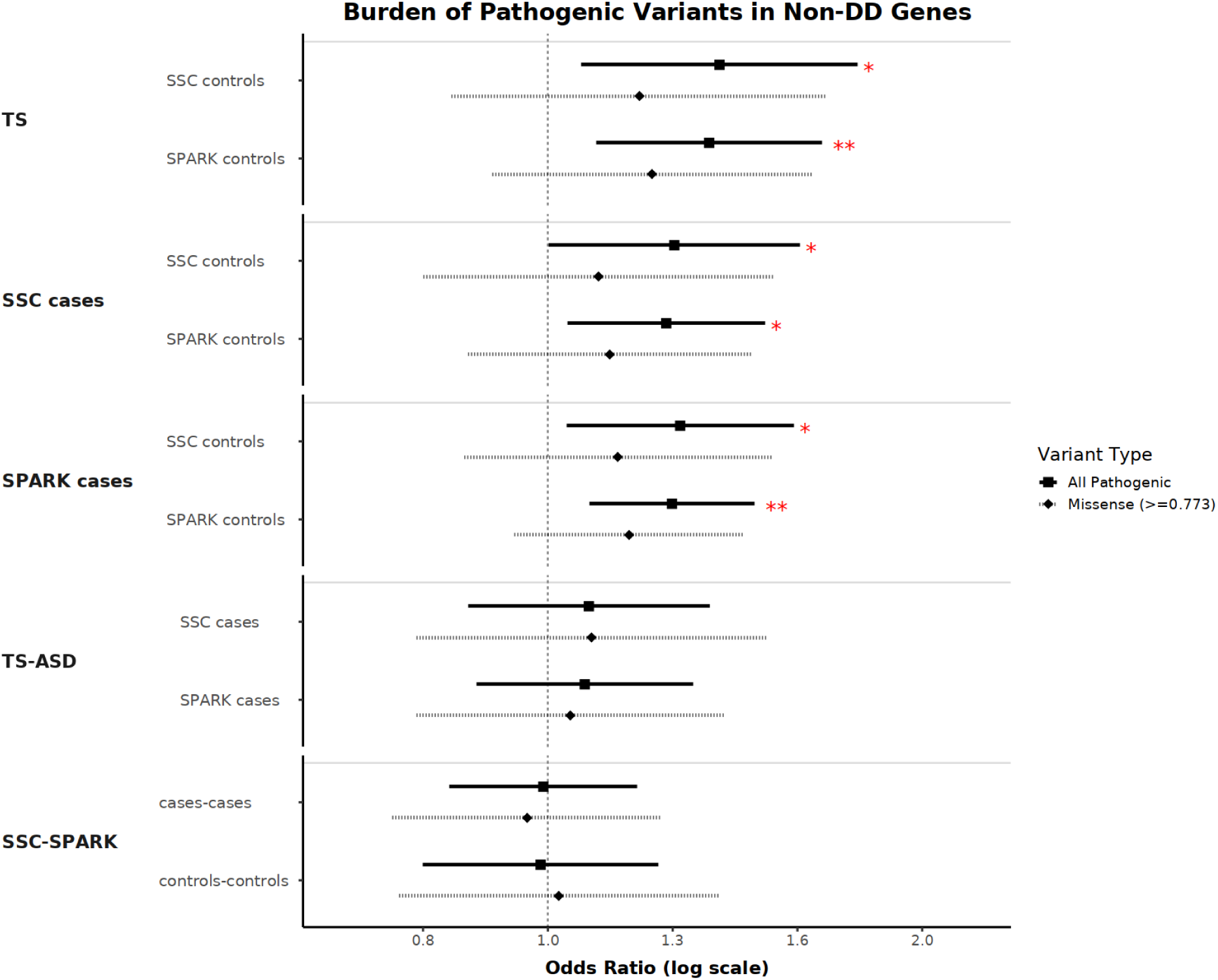
Forest plot of the burden analysis of DNMs between cohorts with the known DD genes removed, split by variant annotations. All pathogenic, PTV + Dmis; Dmis, missense variants with a REVEL score ≥ 0.773.

**Supplementary Figure 12.**
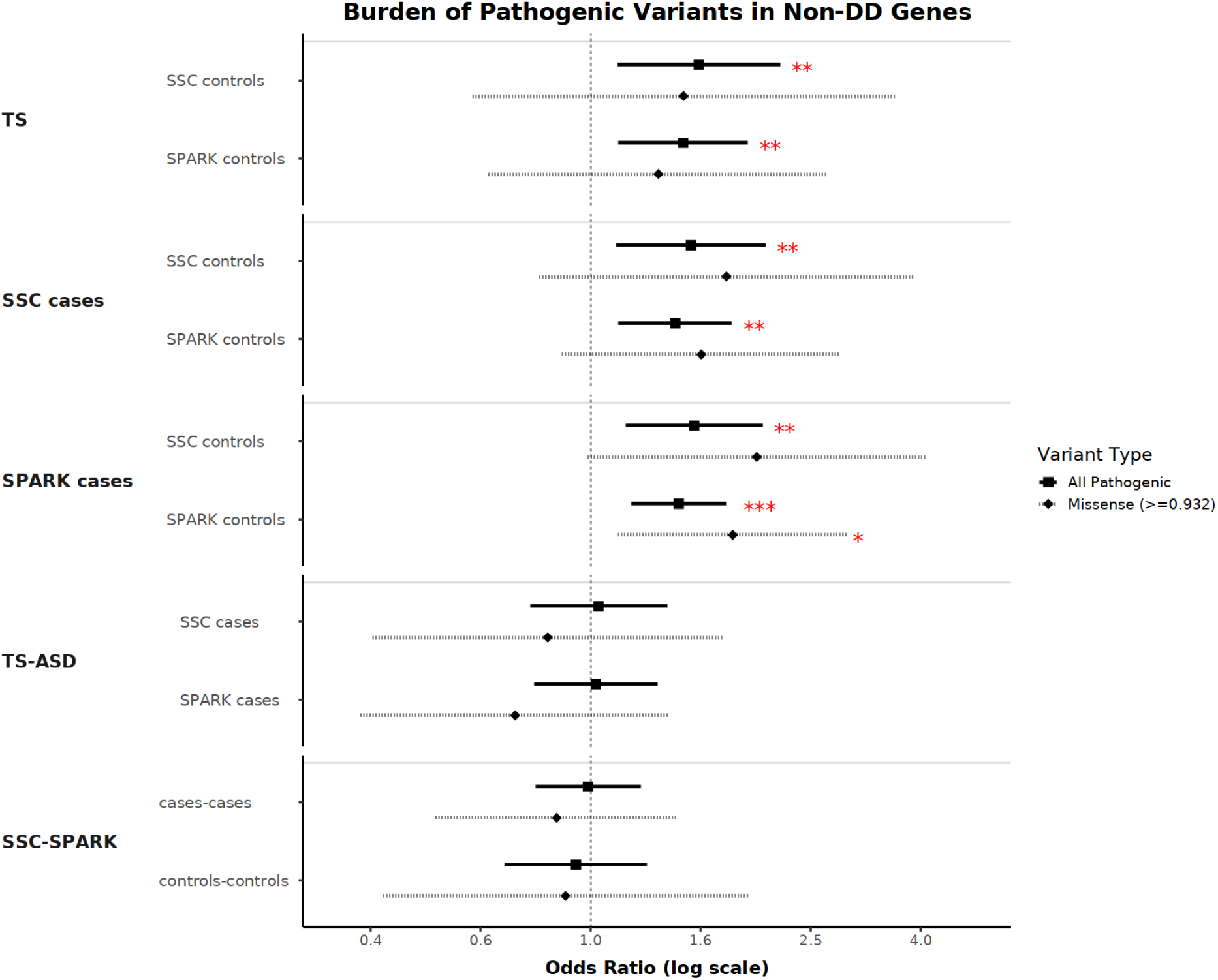
Forest plot of the burden analysis of DNMs between cohorts with the known DD genes removed, split by variant annotations. All pathogenic, PTV + Dmis; Dmis, missense variants with a REVEL score ≥ 0.932.

